# Spatial transcriptomics reveals brain-wide circadian disruption in an Alzheimer’s disease model

**DOI:** 10.64898/2026.01.26.701799

**Authors:** Alon Gelber, Haylie Romero, Dominic Burrows, Daniel S. Whittaker, Daniel Carlin, Eran A. Mukamel, Paula Desplats

## Abstract

Diurnal rhythms in brain transcription align neural, immune, and metabolic processes with the light–dark cycle and are profoundly disrupted in Alzheimer’s disease (AD). However, the regional organization of diurnal transcription in the healthy and diseased brain remains poorly defined. Using large-scale spatial transcriptomics, we mapped 24-hour rhythmic transcription across cortical and subcortical regions of the mouse brain. We identified marked regional differences in rhythmicity, including distinct oscillatory signatures across cortical areas and along the rostro–caudal axis. In the APP23 mouse model of AD, pathology-vulnerable brain regions exhibited early, region-specific disruption of diurnal transcription prior to substantial amyloid plaque deposition. These findings reveal a spatially organized architecture of brain diurnal rhythms and identify early rhythmic dysregulation as a feature of Alzheimer’s disease pathogenesis.

## Main Text

Circadian rhythms are crucial regulators of brain function, rhythmic gene expression, synaptic plasticity, and metabolic cycles (*1*). Precise temporal regulation of these oscillations is essential for processes underlying learning, memory consolidation, and executive function(*1*). Spatial mapping of neural activity across the mouse brain reveals region-specific regulation of the phase and amplitude of circadian rhythms(*2*). Yet, previous studies have not systematically mapped spatially-resolved patterns of rhythmic brain transcription(*3–7*).

Mounting evidence indicates that disruption of intrinsic rhythmicity is a significant driver of cognitive decline and neurodegeneration(*8–10*). Disturbances of daily rhythms are prominent and debilitating symptoms affecting most Alzheimer’s disease (AD) patients, and include dysregulation of sleep/wake cycles(*11*) and exacerbation of cognitive impairment and confusion during the evening (known as “sundowning”)(*12*). Mechanistically, the circadian clock directly influences amyloid-β (Aβ) regulation and plaque formation (*13–15*). Rhythmic DNA methylation of the core clock gene *BMAL1*, (Basic helix-loop-helix ARNT-like protein 1), is altered in AD patient brains and fibroblasts and correlates with altered rhythmic transcription(*16*). These findings support the hypothesis that disruptions to the circadian system can drive neurodegeneration (*9*, *10*).

Despite intensive research, the role of rhythmic gene expression in specific brain regions, and throughout disease progression remains unknown. This gap persists due to the lack of transcriptome-wide, spatially resolved information about the brain-region specific regulation of gene expression in controlled models of AD pathology. Here, we applied spatial transcriptomics (ST) to investigate diurnal gene expression cycles in the mouse brain and to identify alterations in rhythmic expression at early and advanced stages of neurodegeneration in a rodent model of AD(*17–19*). Our data represent a unique resource comprising 65 deeply sequenced mouse brains, covering 23 regions, 4 Zeitgeber time points, and two adult ages in both, non-transgenic controls and APP23-TG mice. We identified profound regional differences in rhythmicity, including distinct oscillatory signatures in the visual cortex compared to somatosensory, motor, and frontal areas. We detected thousands of differentially rhythmic genes whose diurnal expression patterns are disrupted in APP23-TG mice, particularly in brain regions highly vulnerable to AD pathology such as the dentate gyrus and cortex. Notably, rhythmicity was severely disrupted at early disease stages, prior to the overt Aβ pathology. These results suggest that brain-wide disruptions in rhythmic gene expression contribute to AD pathogenesis and may be targeted for clinical intervention.

## Results

### A spatial transcriptomic atlas of rhythmic gene expression across the mouse brain

We used spatial transcriptomics (ST) to comprehensively map rhythmic dynamics of gene expression across multiple brain regions (*18*, *20*). We designed our study using Visium ST to take advantage of the fine-grained spatial resolution (50 µm spot diameter, 100 µm center-to-center spacing) and wide field of view capturing an entire sagittal mouse brain section (Fig. 1A).

**Fig. 1.**
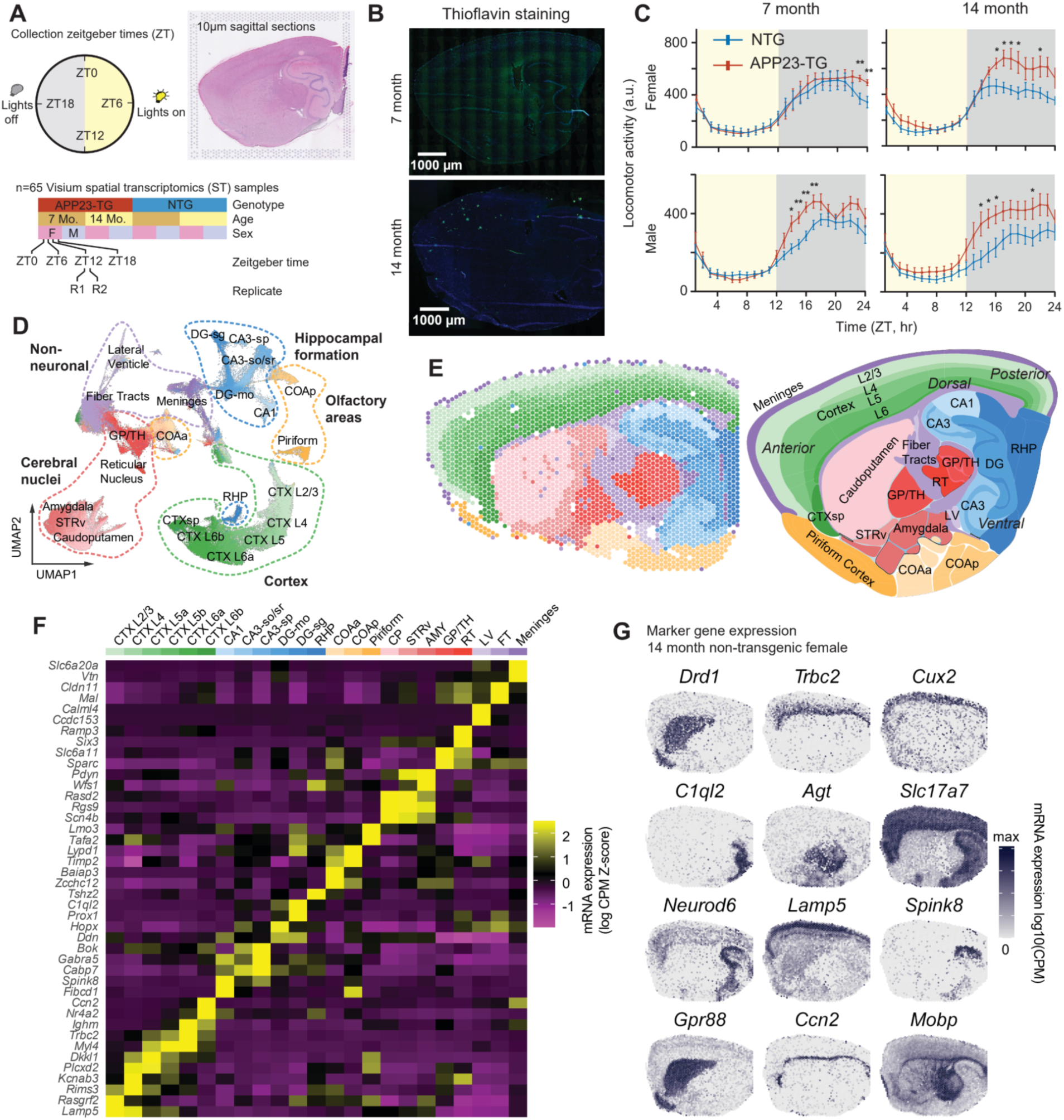
ST analysis of rhythmic gene expression across mouse brain regions and disease stages in the APP23 model of AD. A, Sequential 10-μm sagittal sections of APP23 transgenic (APP23-TG) and nontransgenic (NTG) brains were collected at 7 and 14 months of age, at four time points, including lights on and every six hours after, following monitoring of daily activity rhythms. **B,** The first section was used for ST and the next adjacent sections were used for Thioflavin S staining of plaques. **C,** Mean hourly activity under a 12 h light:12 h dark (LD) cycle at 7 and 14 months; mean ± SEM. Asterisks indicate significant genotype differences at Zeitgeber times (t-test; *p < 0.05, **p < 0.01). **D,** Embedding of 211,899 spots colored by annotated PRECAST clusters (n=65 samples). **E,** Clusters plotted on a representative slice and the Allen reference. **F,** Heatmap showing relative expression of top marker genes for each cluster. Row z-scored Log2 (count per million (CPM) +1). **G,** Examples of spatial expression of marker genes in a 14-month old NTG female.

To detect diurnal rhythms, we maintained mice in a 12:12 light dark cycle and collected brain samples at four Zeitgeber time points (ZT) every six hours from the lights on time (ZT0) (Fig. 1A). We applied this paradigm to male and female APP23 transgenic(*21*) (APP23-TG) and non-transgenic (NTG) littermate control mice at two key ages: 7 months, representing early-symptomatic mice, and 14 months, when transgenic mice show significant amyloid plaque burden. From each mouse, we collected a 10 µm thick section for ST and an adjacent 10 µm thick section that we stained for Thioflavin S to identify Aβ plaques (Fig. 1B). This data resource represents an unprecedented, comprehensive spatial map of brain diurnal gene expression.

The spatial resolution of the ST platform enables capturing mRNA from 1-10 cells, depending on the local cell density. We used spatially-aware clustering to define groups of ST spots based on similar gene expression and spatial contiguity(*22*). We chose cluster analysis parameters to match the anatomic and transcriptomic resolution of the Allen Brain Mouse Reference Atlas at the subclass level(*23*).

Our ST data provide comprehensive coverage of the mouse brain transcriptome at 211,899 total ST spots with an average of 15,696 unique transcripts (4,126 unique genes) detected per spot. We grouped these data into 23 anatomically and molecularly defined clusters (Fig. 1D-F). Cortical layers were separated based on layer-specific excitatory neuron markers, as were the major hippocampal subfields and layers (CA3 stratum radiatum (sr) and stratum pyramidale (sp); CA1; dentate gyrus (DG) molecular layer (mo) and stratum granulare (sg)) and retrohippocampus. We also identified major subcortical clusters including anterior and posterior cortical amygdala (marked by *Baiap3, Zcchc12* and *Lypd1*)(*24*), basal ganglia (cerebral nuclei) subregions such as reticular nucleus (*Ramp3)* and globus pallidus (*Sparc*), and piriform cortex (*Lmo3*, *Tafa2*). Finally, we identified clusters corresponding to white matter (*Mal*), ventricles (*Calml4*), and/or meninges (*Vtn)*. We annotated each cluster based on the expression of marker genes (Fig. 1F,G).

### Brain-region-specific rhythms of gene expression

Our experimental design enables testing rhythmic gene expression in the context of multiple variabless, including sex, age and genotype. We developed a flexible framework to detect 24-hour rhythmicity of gene expression using negative binomial harmonic regression of pseudobulk profiles with DESeq2(*25*). The model includes a rhythmic component expressed as a sinusoidal function of zeitgeber time (ZT). For each gene, we tested the significance of diurnal rhythmicity using a likelihood ratio test against a reduced model with no dependence on ZT. We validated that our approach is consistent with widely-used methods from the Metacycle package(*26*), while ours accommodates key covariates (Fig. S1A-D).

The circadian clock is defined by a conserved core oscillating circuit comprising *Per1, Per2, Bmal1 (Arntl)*, *Dbp, Cry1* and *Nr1d2*(*27*). This circuit oscillates autonomously with a ∼24-hour period in most tissues, including the brain, and is entrained to the ambient day-night cycle via light-activated neurons in the supra-chiasmatic nucleus (SCN) of the hypothalamus(*28*). The core molecular clock regulates a range of tissue- and cell type-specific effector genes in sync with the diurnal cycle to optimize cellular physiology to tissue-specific requirements. In the mouse brain, we found that core clock genes were significantly rhythmic in most brain regions (FDR<0.05, relative amplitude>0.05; Fig. 2A). The amplitude and phase of each core clock gene, defined by the ZT of peak expression (acrophase), were consistent across the brain (Fig. 2A,B). We found 119 rhythmic genes (FDR<0.2) in at least 10 out of 23 brain regions, and 52 genes were rhythmic (FDR<0.05) in ≥15 regions (Fig. 2B, Supplementary Table S1).

**Fig. 2.**
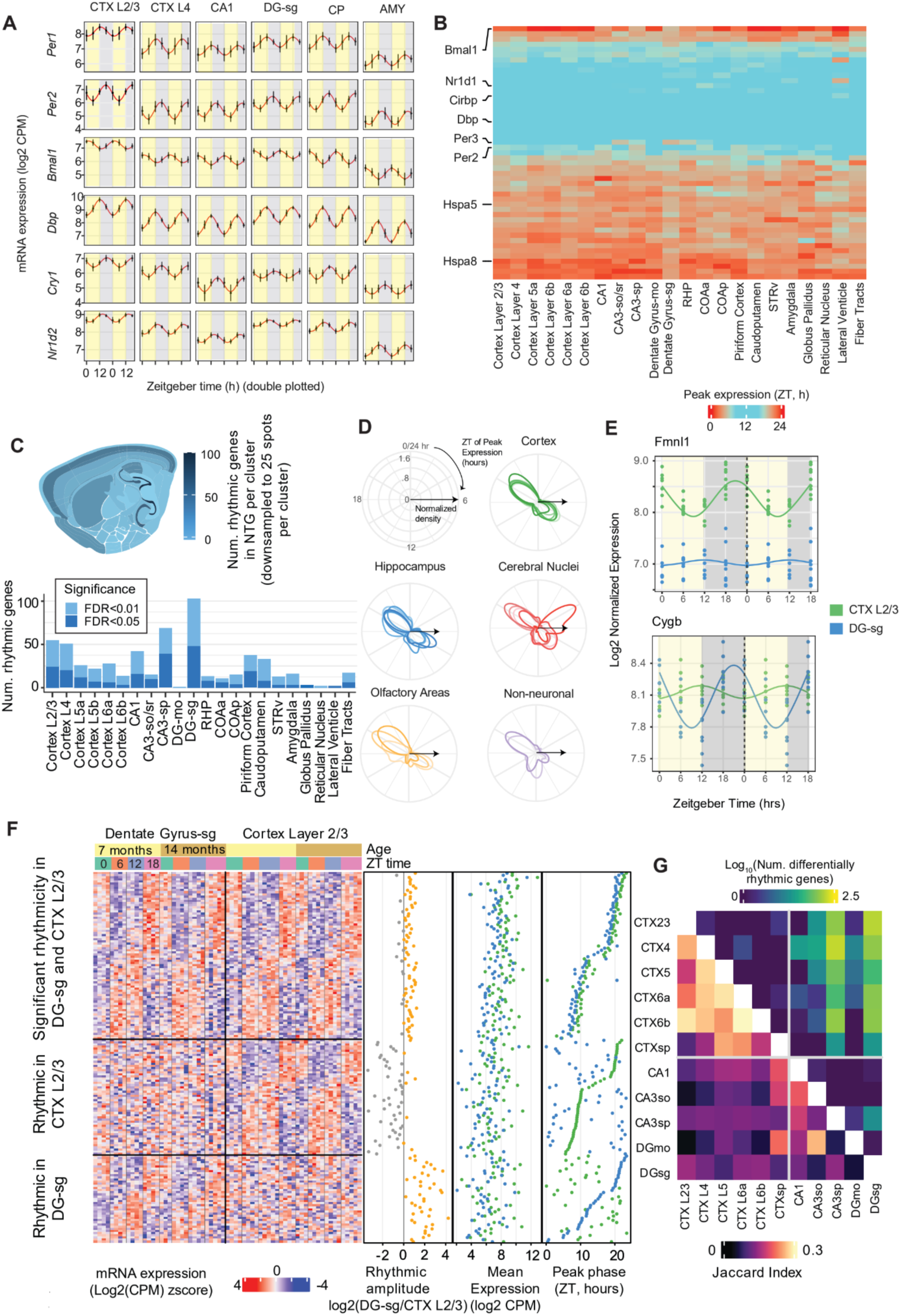
Spatial transcriptomic profiling of 24-hour rhythmicity in gene expression across the brain. A, Expression of core clock genes across brain regions. Data are double-plotted with sinusoidal model fit (red). Points and error bars show mean ± SD. **B,** Consistent phase of expression for the top rhythmic genes shared across brain regions (FDR<0.1 in at least 15 regions). **C,** Number of significantly rhythmic genes in each brain region in NTG mice (FDR<0.05, likelihood ratio test). Each cluster was downsampled to 25 spots. **D,** Polar plots of the distribution of peak phases for rhythmic genes. **E,** Region-specific rhythmic genes in cortex layer 2/3 (*Fmnl1*, green) and the dentate gyrus (*Cygb*, blue). **F,** Expression of rhythmic genes shared between cortex layer 2/3 and the dentate gyrus and region-specific rhythmic genes (DRGs, FDR<0.1 likelihood ratio test). **G,** Overlap of rhythmic genes and number of DRGs (LRT FDR<0.1) for all pairs of hippocampal and cortical clusters.

The number of significantly rhythmic genes varied across brain regions (FDR<0.05, Supplementary Table S2). To compare brain regions without bias due to differences in size, we downsampled the data from each region to 25 randomly selected ST spots per sample (Fig. 2C). Hippocampal and cortical regions had the largest numbers of significantly rhythmic genes (FDR<0.05). The granule cell layer of the dentate gyrus (DG-sg) had the most rhythmic genes (142 genes downsampled, 552 full dataset), followed by CA3sp (101 genes downsampled, 357 full dataset) and CA1 (57 genes downsampled, 172 full dataset) of the hippocampus. The cortex was also highly rhythmic, especially the upper layer 2/3 (63 genes downsampled, 1047 full dataset). Deep cortical layers, piriform cortex, and the caudoputamen had a moderate number of rhythmic genes, while cerebral nuclei and non-neuronal areas had the fewest (Fig. S2A).

The peak phase of rhythmic genes is largely consistent across tissues in humans(*29*), but differences in phase between brain regions have not been characterized. We found that the phase of rhythmic gene expression was bimodally distributed in the mouse brain, with most genes peaking around ZT8 or ZT20 hours (Fig. 2D). An exception was the reticular nucleus of the thalamus (RT), where 46 rhythmic genes peaked around ZT4. The RT serves as a relay between thalamic and cortical circuits and plays a critical role in sleep maintenance(*30*). Because non-rapid eye movement (NREM) sleep patterns are driven by reciprocal thalamic-cortical coupling, the unique phase profile of the RT may reflect its role in coordinating sleep stages(*31*). This finding highlights the importance of spatially resolved analysis of rhythmic gene expression, particularly for small but critical structures like the RT.

Our data allowed us to identify genes that have a brain-region specific rhythmic expression profile, potentially due to regional differences in diurnal activity(*3*). We found examples of genes with similar mesor levels across regions, but showing significantly different amplitude and/or phase of diurnal rhythmicity. To illustrate, we compared two regions with the highest number of rhythmic genes (DG-sg and Cortex L2/3, Fig. 2F). These regions shared 232 rhythmic genes (FDR<0.1), but each also had specific sets of rhythmic genes in only one region (1051 genes in L2/3, 419 in DG-sg). For example, the cell structure regulator Formin-like 1 (*Fmnl1*) had a strong diurnal rhythm in Cortex L2/3, but no apparent rhythmicity in DG-sg (Fig. 2E). By contrast, the oxygen regulator cytoglobin (Cygb) was strongly rhythmic in DG-sg but not Cortex L2/3.

To assess differential rhythmicity between regions, we used a likelihood ratio test that compares a model with independent harmonic coefficients in each region to a null model with shared harmonic components (FDR<0.1, Supplementary Table S3). This test is sensitive to differences in either phase or amplitude. We found that the rhythmic transcriptome is relatively conserved across cortical layers, with a strongly overlapping set of rhythmic genes (Jaccard index=0.21-0.28) and only a handful differentially rhythmic genes (DRGs, Fig. 2G, Supplementary Fig. S2D). By contrast, hippocampal clusters shared fewer rhythmic genes and had substantial differential rhythmicity with cortical regions.

### Regional specialization of rhythmic gene expression across major spatial axes of the hippocampus and cortex

Our ST dataset not only reveals distinct rhythmic regulation across major brain structures (Fig. 1E), but also resolves spatial differences within key structures. We leveraged this spatial resolution to compare the dorsal and ventral poles of the hippocampus (Fig. 3A) and cortical regions ranging from rostral motor and somatosensory areas to caudal visual cortex (Fig. 3G).

**Fig. 3.**
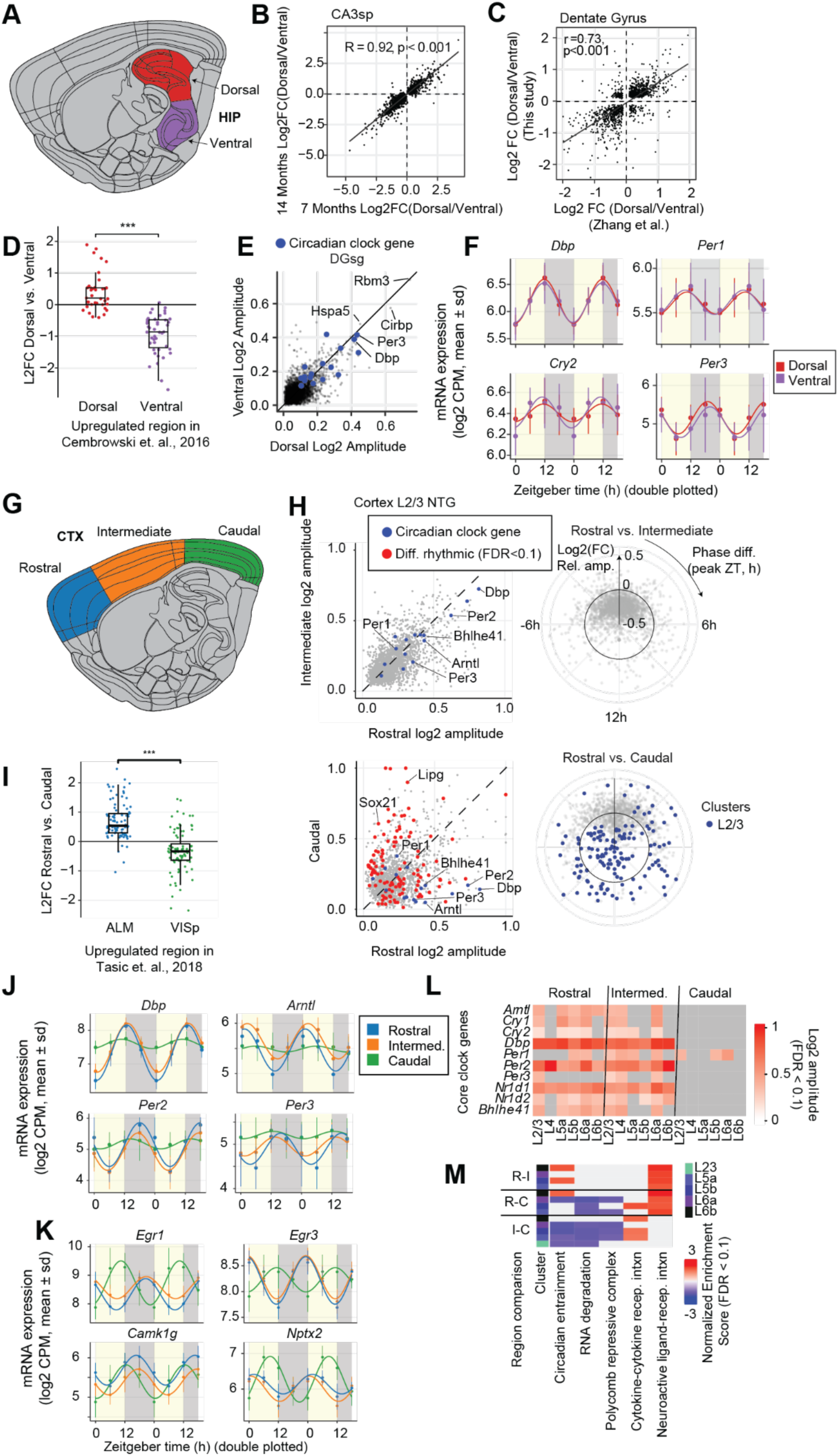
Dorso-ventral transcriptional gradients and subregion-specific rhythmicity across hippocampus and cortex. A, Hippocampal subregions**. B,** Dorsal/ventral log2 fold changes in representative hippocampal cluster in NTG mice at 7 versus 14 months (spearman r = 0.92, p < 0.001) **C,** Dorsal/ventral log2 fold changes in dentate gyrus comparing this dataset to Zhang et al. 2018 (*32*) bulk RNA-seq (r = 0.73, p < 0.001) **D,** Boxplot of dorsal vs. ventral log2 fold changes for genes annotated as dorsal or ventral markers in Hipposeq (Cembrowski et al. 2016(*35*)); points are individual genes (two-tailed t-test, ***p < 0.001). **E,** Scatter of log2 amplitudes in dorsal vs. ventral DG (FDR < 0.1); core clock genes highlighted. **F,** Rhythmic expression of representative transcripts in DG: points, mean ± SD at each ZT; curves, harmonic regression fits (dorsal, red; ventral, purple).**G,** Cortical subregions. **H,** Scatter of log2 rhythmic amplitudes in cortical layer 2/3 for rostral versus intermediate cortex (differential rhythmicity FDR < 0.1 in red); core clock (blue) genes highlighted. Inset polar plot depicts phase and amplitude differences. **I,** Boxplot of rostral vs. caudal log2 fold changes for genes annotated as ALM- or VISp-enriched (Tasic et al. 2018(*34*)); points are individual genes (two-tailed t-test, ***p < 0.001). **J,** Rhythmic expression of core clock genes across cortex (average over all layers): mean ± SD with harmonic regression fits for differentially rhythmic genes. **K,** Rhythmic expression for phase-shifted genes across cortex subregions; mean ± SD and harmonic fits. **L ,** Heatmap of log2 rhythmic amplitude for core clock genes (rhythmicity FDR < 0.1) across layers L2/3, L5a, L5b, L6a, L6b within the rostral, intermediate and caudal cortex. **M,** Heatmap of functional enrichment scores using GSEA for gene expression differences between cortical regions (R-I, rostral versus intermediate; R-C, rostral versus caudal; I-C, intermediate versus caudal) (FDR < 0.1).

Thousands of genes showed distinct expression patterns between the dorsal and ventral poles of hippocampal areas, with the most DEGs observed in area CA3 (Fig. S3D, Supplementary Table S4). We confirmed previous findings of a robust dorsal-ventral gradient of expression of multiple marker genes, including patterning factors like *Epha7* and *Nr2f2* (Fig. S3D), as well as *Trhr* and *Lct* (Fig. S3E-F)(*32*, *33*). The dorsal-ventral gradient of gene expression was consistent with previous studies using bulk RNA-seq from the dorsal and ventral poles of the DG(*32*) (Fig. 3C, Pearson r=0.73, p<0.001). We found that dorsal genes in all 4 hippocampal clusters were functional enriched for long-term potentiation (FDR<0.05, see Methods). Ventral-enriched genes had more variable functional profiles (Fig. S3H).

Despite the large transcriptional differences in mean expression between dorsal and ventral hippocampal regions, we found highly consistent rhythmic modulation in both poles of the hippocampus (Fig. 3H). Core clock genes showed nearly identical amplitude and phase of expression (Fig. 3H). No genes had significantly different rhythmicity in dorsal vs. ventral hippocampus (interaction between region and ZT, FDR<0.1). Thus, although dorsal and ventral hippocampal regions have robust differences in gene expression, their circadian regulation appears to be highly consistent.

The neocortex is organized along the rostro-caudal axis in major regions specialized for cognitive functions, from the primary visual cortex (VISp) at the caudal pole to somatosensory in middle (intermediate) and motor and frontal cortices at the rostral end. Single cell transcriptomics has identified hundreds of genes that are differentially expressed in VISp vs. antero-lateral motor (ALM) cortex(*34*). We found thousands of DE genes in pairwise comparisons of rostral vs. caudal, rostral vs. intermediate, and intermediate vs. caudal regions (Fig. 3H, FDR<0.05, Supplementary Table S5). Rostral vs. caudal DEGs were largely consistent with previous scRNA-seq from VISp and ALM(*34*) (Fig. 3I).

Caudal genes were significantly enriched in functional categories including “circadian entrainment” and “RNA degradation” (Fig. 3M). These genes included components of intracellular calcium signaling (*Adcyap1, Camk2d, Cacna1c/d/h*), MAPK signaling (*Mapk3*), activity-regulated genes (*Fos, Rasd1*), and other modulators of circadian input pathways (*Adcy8, Grin2b, Prkg2, Prkacb*) (Fig. 3M). Importantly, this analysis did not explicitly account for ZT and thus reflects regional differences in the mesor, or mean gene expression across the day. Our data thus suggest the caudal cortex, despite having less rhythmic expression of core clock genes, may be transcriptionally primed for circadian entrainment and activity-dependent signaling.

Furthermore, analysis of rhythmic transcription identified robust differences in gene expression in the caudal cortex compared to rostral and intermediate regions. There were up to 390 DRGs between caudal and rostral or intermediate cortex across cortical layers, while ≤3 genes were significant DRGs between rostral and intermediate cortical regions (Fig. 3H, FDR<0.1, Supplementary Table S6). This was notable, since there were hundreds of genes with significantly different expression mesor in rostral, intermediate and caudal cortical regions (Supplementary Fig. S3I).

Notably, the rhythmicity of several core clock components was attenuated or abolished in the caudal cortex. These included *Arntl (Bmal1), Nr1d1 (Rev-Erbα), Nr1d2 (Rev-Erbβ), Dbp, Per2,* and *Bhlhe41 (DEC2)*, which were substantially more rhythmic in rostral and intermediate compared with caudal cortex (Fig. 3J). The rhythmic amplitude of core clock genes was consistently lower in caudal regions across cortical layers (Fig. 3L). Gene ontology analysis showed that 177 DRGs with stronger rhythmicity in rostral compared with caudal cortex were enriched for circadian rhythm function (FDR<0.001, Supplementary Table S7).

We further identified a distinct set of 169 genes that were rhythmic in caudal cortex, but lacked significant rhythmicity in rostral or intermediate regions. These caudal-specific rhythmic genes were enriched for regulators of MAPK signaling, such as *Dusp5, Dusp6, Adcy8,* and *Elk1* (Fig. 3H). The robust diurnal rhythmicity of these genes shows that caudal cortex does not lack a 24-hour clock, despite the reduced rhythmicity of core clock components.

We identified 22 genes that had significant rhythmic expression in both caudal and rostral/intermediate cortex but with significant differences in phase, including *Egr1, Egr3, Camk1g*, and *Nptx2* (Fig. 3L). The phase shift between rostral vs. caudal regions was consistent across cortical layers. These genes are involved in activity-dependent transcriptional programs, synaptic signaling, and calcium-mediated pathways, and their coordinated phase shifts across layers suggest region-specific regulation of their rhythmic expression. For example, in nocturnally active species like mice, visual stimulation during the light phase could drive increased expression of neural activity-related genes in caudal cortex, whereas somatosensory and motor regions would be more active during the dark phase. These findings are consistent with a recent neural activity mapping by c-Fos protein expression, which also identified an opposite phase of circadian rhythmicity in visual cortex compared to other cortical regions(*2*). Finally, we confirmed that the within-region expression gradients and specific rhythmic patterns we observed in NTG animals were also present in APP23-TG mice, and thus seems spared from pathology-associated alterations (Supplementary Fig. S4).

### Disruptions of brain-region specific transcriptomes in APP23-TG AD mice

To investigate the impact of AD-related pathology on rhythmic gene expression across the brain, we performed ST in APP23-TG mice(*9*, *21*, *36*). Male and female APP23-TG mice had disrupted circadian behavioral patterns, fragmented sleep and increased locomotor activity in the home cage during the dark phase(*9*) (cage activity) (Supp. Fig. S5A,B), consistent with our previous report(*9*). APP23-TG mice show early circadian disruption at 7 months of age, despite the absence of overt amyloid pathology, with a longer intrinsic circadian period (free-running period) than NTG mice, measured in constant darkness, (p ≤ 0.05, Supplementary Fig. S5D). By contrast, at 14 months of age both APP23-TG and NTG animals had equivalent intrinsic circadian periods, which were longer than that of 7 month NTG animals.

Our findings are in line with the characterization of APP23-TG mice showing development of progressive neuropathology starting around 6 months of age with sparse Aβ buildup, followed by accumulation of plaques at 7-9 months in the cortex which then spread to the hippocampus(*37*) (Fig. 1B, Supplementary Fig. S6A,B). This was accompanied by increased expression of reactive astrocyte marker, GFAP (Supplementary Fig. S6C,D). As the disease progressed, both Aβ plaque burden and astrogliosis increased as did the development of microgliosis throughout the cortex (Supplementary Fig. S6E-G)(*21*).

To determine whether AD pathology impacts brain region-specific gene expression, we identified differentially expressed genes (DEGs) between APP23-TG and NTG mice using conservative, statistically rigorous criteria (see Methods)(*25*, *38*). We observed up to 498 DEGs per brain region, with the largest changes located in the hippocampus, dentate gyrus and cortex (FDR<0.1, Fig. 4A, Supplementary Table S8). In the cortex, the greatest number of DEGs was found in layer 5b in both the pre-amyloid (7 month old) and advanced pathology (14 month) groups (Fig. 4B). The distribution of DEGs across layers mirrored the Aβ plaque burden in 14-month old APP23-TG mice, as quantified by the proportion of plaque-associated spots per cortical layer (Fig. 4C).

**Fig. 4.**
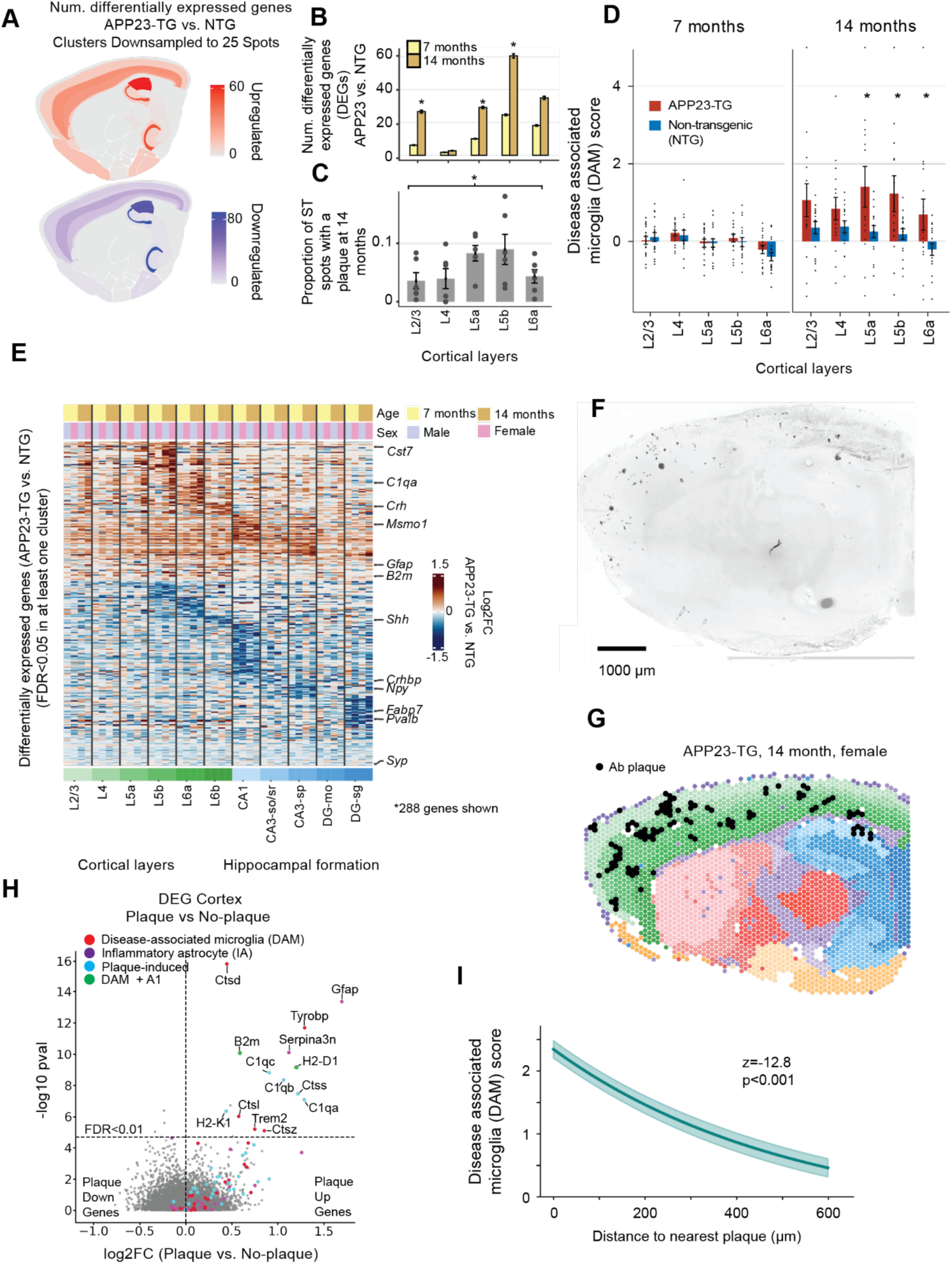
Altered spatial patterns of mRNA expression in APP23-TG mice. A, Number of differentially expressed genes (DEGs) in APP23-TG vs. NTG mice in each brain region (FDR<0.05) at 14 months. Clusters were downsampled to 25 spots per sample. **B,** Number of DEGs in APP23-TG mice per cortical layer at 7 and 14 months. DE analysis was performed by downsampling cortical layers to 100 spots 20 times and computing the mean and standard error of the number of DEGs with a Wald test (FDR<0.05). *: p<0.05 (Wilcoxon test, 7 vs. 14 months). **C,** Mean Aβ plaques per ST spot in each cortical layer in 14 month APP23-TG mice (errorbar: SEM; dots: individual samples). *: p<0.05, mixed model controlling for sex and sample as a random effect (p<0.05). **D,** Disease associated microglia (DAM) score for cortical layers (errorbar: SEM; dots: individual samples). *: p<0.05, T-test, APP23-TG vs NTG. **E,** 288 DEGs in cortex and hippocampus (FDR<0.05, LRT: ∼sex+age+genotype vs. ∼sex+age; mean(|L2FC|)>0.3, max(|L2FC|)>0.7). **F,** Example of Thioflavin S staining in 10 µm sections adjacent to sections used for ST. **G,** Example 14 month APP23-TG sample with annotated plaques. **H,** 88 DEGs between Aβ plaque-associated spots and non-plaque associated spots in the cortex (FDR<0.1, LRT: ∼plaque+(1|sample) vs. (1|sample) ). Significantly DE DAM, inflammatory astrocyte (A1) and plaque-induced (PI) genes(*43*) are highlighted. **I,** Exponential fit of the average DAM score for each spot as a function of the distance to the nearest plaque. *908 genes with FDR<0.1 (LRT: ∼plaque_distance+(1|sample) vs. (1|sample) ).

In the cortex and hippocampus, DEGs included hallmarks of neuroinflammation in AD such as microglial genes *Cst7* and *B2m* (Fig. 4E). Neurodegeneration has been linked to an increased proportion of disease-associated microglia (DAM)(*39*). We created a gene expression-based DAM score using known markers of Alzheimer’s DAM(*39–42*). The DAM score was significantly higher in 14 month old APP23-TG mice compared with NTG controls, particularly in deep cortical layers (p<0.05, Fig. 4D).

### Genes associated with AD pathology are activated in the proximity of Aβ plaques in APP23-TG mice

Aβ plaques can alter the regulation of nearby cells by inducing expression of genes involved in myelination, complement signaling, oxidative stress and inflammation(*43*). To address the spatial relationship between Aβ plaques and altered gene expression in our ST data, we used thioflavin S staining to identify plaque associated spots in brain sections immediately adjacent to those used for ST (Fig. S6C). The average number of Aβ plaques per sample increased from 7±6 at 7 months of age to 42±37 at 14 months in APP23-TG animals (mean ± SD, p<0.01, n=9; Fig. S6B).

To identify plaque-induced genes, we analyzed DEGs in plaque-associated vs. non-plaque-associated ST spots in the cortex using a mixed effects model to account for inter-sample variability. We found 47 upregulated and 41 downregulated DEGs in plaque-associated spots (FDR<0.1, Supplementary Table S9). The DEGs included known markers of disease pathology including DAM-associated, inflammatory astrocyte (A1) and other plaque-induced genes like *C1q(a-c)*, *Ctss* and *Npc2* (Fig. 4H). Notably, the plaque-induced genes overlapped with findings from another AD mouse model, App^NL-G-F^ (22 shared genes; p<0.0001 Fisher’s exact test)(*43*), validating the findings. Furthermore, we found that 11 key DAM genes had higher expression in spots neighboring a plaque compared with non-plaque associated spots (FDR<0.01, Fig. 4H). The DAM score was significantly higher in spots located <200 µm from a plaque compared to spots with no nearby Aβ plaques (Fig. 4I).

### Increased rhythmic amplitude of gene expression in APP23-TG mice

Rhythmic gene expression is disrupted in the AD brain(*8–10*, *47*, *48*), but the regional pattern of disruption has not been established. We identified hundreds of genes that gained or lost rhythmicity in APP23-TG compared to NTG animals. Notably, we observed a greater number of rhythmic genes in APP23-TG animals across all brain regions compared to controls (Fig. 5B), and an increase in amplitude in most rhythmic genes brain-wide (Fig. 5A). To statistically test the difference in rhythmicity for individual genes, we used a conservative, permutation-based procedure to generate an empirical null distribution (See Methods, Fig. 5C). The increased rhythmic amplitude in APP23-TG mice was most pronounced in 7 month old animals, though it was still evident in 14 month old mice with more advanced pathology (Fig. 5C). Consistent with this global increase in rhythmic amplitude, hundreds of genes with minimal or undetectable rhythms in NTG controls acquired clear 24 h oscillations in 7-month-old APP23-TG mice. In the dentate gyrus, *Tet3*, a methylcytosine dioxygenase that initiates DNA demethylation(*49*), and *Cxxc5*, which antagonizes Wnt/β-catenin signaling and has been linked to AD(*50*), both became significantly rhythmic in APP23-TG (interaction LRT FDR < 0.05; Fig. 5E). In cortex layer 2/3, APP23-TG induces robust diurnal oscillations in ubiquitin regulators (*Midn*, *Otud1*, *Uxt*)(*51*) and phosphatases (*Dusp4*, *Dusp6*), with peak expression clustering around the transition from dark to light. The increased rhythmicity of protein quality control and signal-termination pathways in cortex may represent an adaptation to early pathological changes (interaction LRT FDR < 0.05; Fig. 5F).

**Fig. 5.**
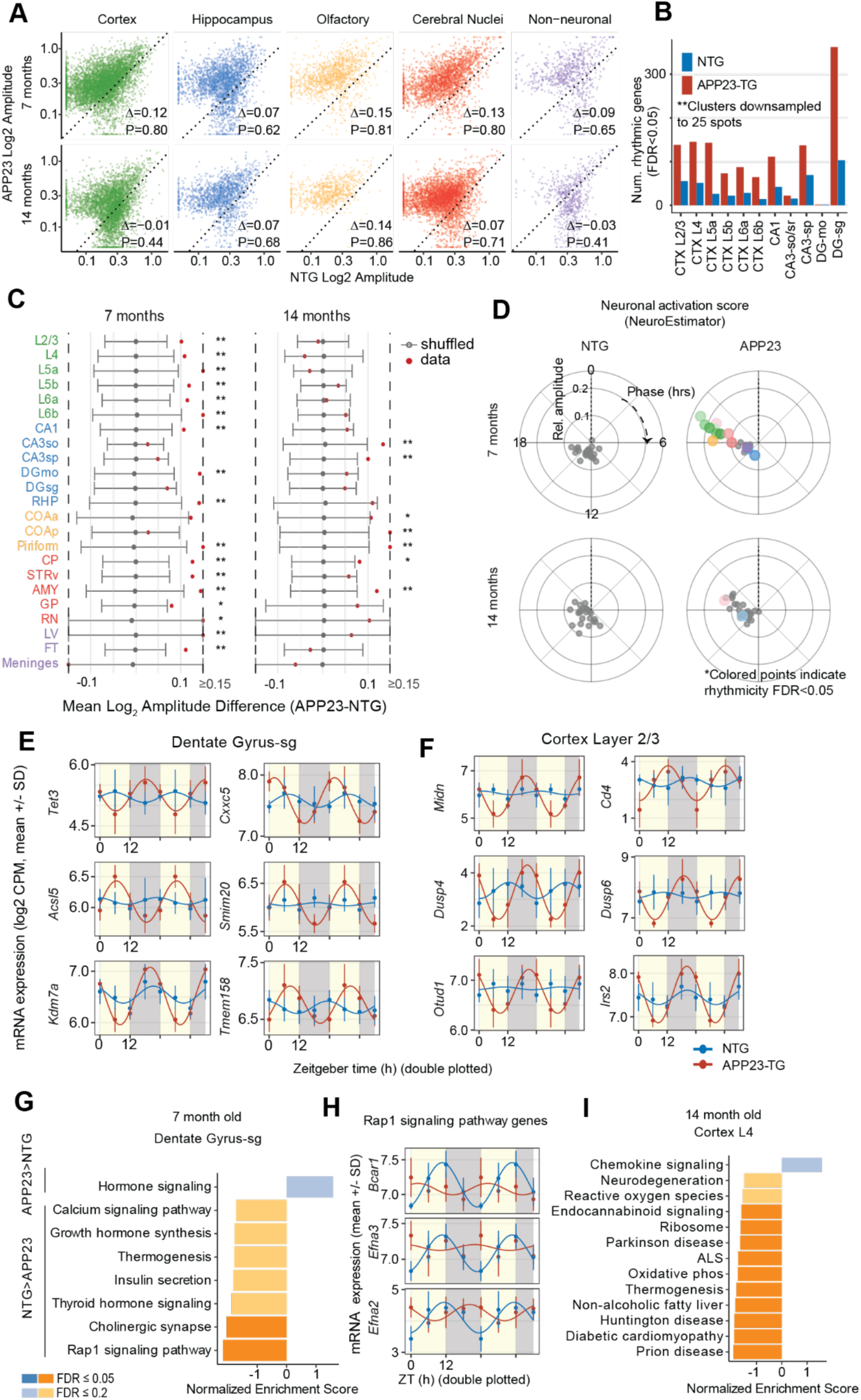
Alterations in diurnal rhythmicity of gene expression in APP23-Tg mouse brain. A, Log2 amplitude of rhythmic expression in APP23 vs. NTG (dashed line shows slope=1). **B,** Number of significantly rhythmic genes in APP23-TG and NTG (FDR<0.05) across cortical and hippocampal regions. **C,** Difference in mean amplitude of rhythmicity between APP23 and NTG for each cluster (red points) plotted on null distribution of the same metric from 500 random permutations of the genotype labels. Grey error bars show 95% confidence interval of the empirical null distribution from 500 random permutations of the genotype labels (*, FDR<0.05). **D,** Phase and amplitude of mean neuronal activation score(*44*) by cluster, age and genotype. Significantly rhythmic clusters (LRT FDR<0.05) are indicated by the colored points while non-rhythmic scores are gray. **E,** Expression of several example genes with greater rhythmicity in APP23-TG compared to NTG (interaction LRT FDR<0.05) in dentate gyrus. **F,** Same as previous in cortex layer 2/3 . **G,** Gene set enrichment analysis on log2 amplitude difference for rhythmic genes in dentate gyrus in 7 month-old animals(*45*, *46*).**H,** Expression of three Rap1 signaling genes (Bcar1, Efna3, Efna2) in cortex layer 4 at 14 months with diminished rhythmicity in APP23-TG compared to NTG. **I,** Gene set enrichment analysis on log2 amplitude difference for rhythmic genes in cortex layer 4 in 14 month-old animals(*45*, *46*).

We found significant functional enrichment among genes with decreased rhythmic amplitude (Fig. 5G). Genes losing rhythmicity in dentate gyrus of 7-month-old APP23-TG animals, including *Bcar1, Efna2,* and *Efna3,* were enriched for pathways linked to AD, like Rap1 signaling, cholinergic synapse, insulin metabolism, thermogenesis, and calcium signaling, (FDR<0.05, Fig.5G,H) (*52*, *53*)(*54*, *55*). On the other hand, genes that gained rhythmicity in APP23-TG were enriched for insulin secretion (e.g. *Irs2*) and parathyroid hormone signaling in cortex layer 2/3.

### Activity dependent genes gain rhythmicity at early stages of AD pathology

Daily rhythms in neural activity lead to oscillation of a set of immediate early genes (IEGs), as recently shown by immunostaining of c-Fos protein(*2*). We found that the genes that gained rhythmicity in APP23-TG cortex and dentate gyrus were enriched in neuronal activity-induced genes (FDR<0.05)(*56*). Because IEGs are rapidly induced by neural activity, shifts in their diurnal phase and/or amplitude suggest altered daily rhythms of cortical and hippocampal neural activity in APP23-TG.

To investigate this, we estimated neural activity scores based on the combined expression of a panel of activity-dependent genes(*44*). We compared these scores between APP23-TG and NTG mice at 7 and 14 months across brain regions using a linear mixed-effects model. We found no significant differences in the mesor of neural activity scores between the genotypes. Activity scores for 12 of the 23 clusters had a significant 24-h rhythm in 7-month-old APP23-TG mice, and in 2 clusters in 14-month-old APP23-TG mice (FDR<0.05, Fig. 5D). By contrast, NTG mice had no significant rhythmicity in neural activity scores (Fig. 5D). Using a likelihood ratio test, we established that APP23-TG mice had significantly higher rhythmicity of the activity score than NTG controls in all cortical clusters and in the DG at 7 months (FDR<0.05). These findings suggest early amyloid pathology may drive synchronized and phase-shifted neuronal activation across the brain.

### Brain region-specific phase shifts in APP23-TG mice

Differential rhythmicity testing revealed widespread shifts in the phase and/or amplitude of daily gene expression rhythms in APP23-TG mice relative to NTG controls at 7 months of age (Fig. 6A, Supplementary Table S10). These shifts were especially pronounced in cortex and in the subgranular zone of the dentate gyrus of the hippocampus, and included both phase advancement and delay. While some genes were only modestly shifted, others had opposite phase (12 h shift) in APP23-TG mice, such as *Nr4a1*, *Dusp1*, and *Trib1*. Notably, core clock genes *Per1* and *Per2*, *Cry2*, and *Bhlhe40* (DEC1) were phase-delayed by ∼6 h in APP23-TG mice compared to NTG (Fig. 6B,C). Consistent with this shift in the core molecular clock, a cortex-wide phase map revealed that many DRGs in cortical clusters had similar phase delays. By contrast, phase advances were more variable across cortical layers (Fig. 6G).

**Fig. 6.**
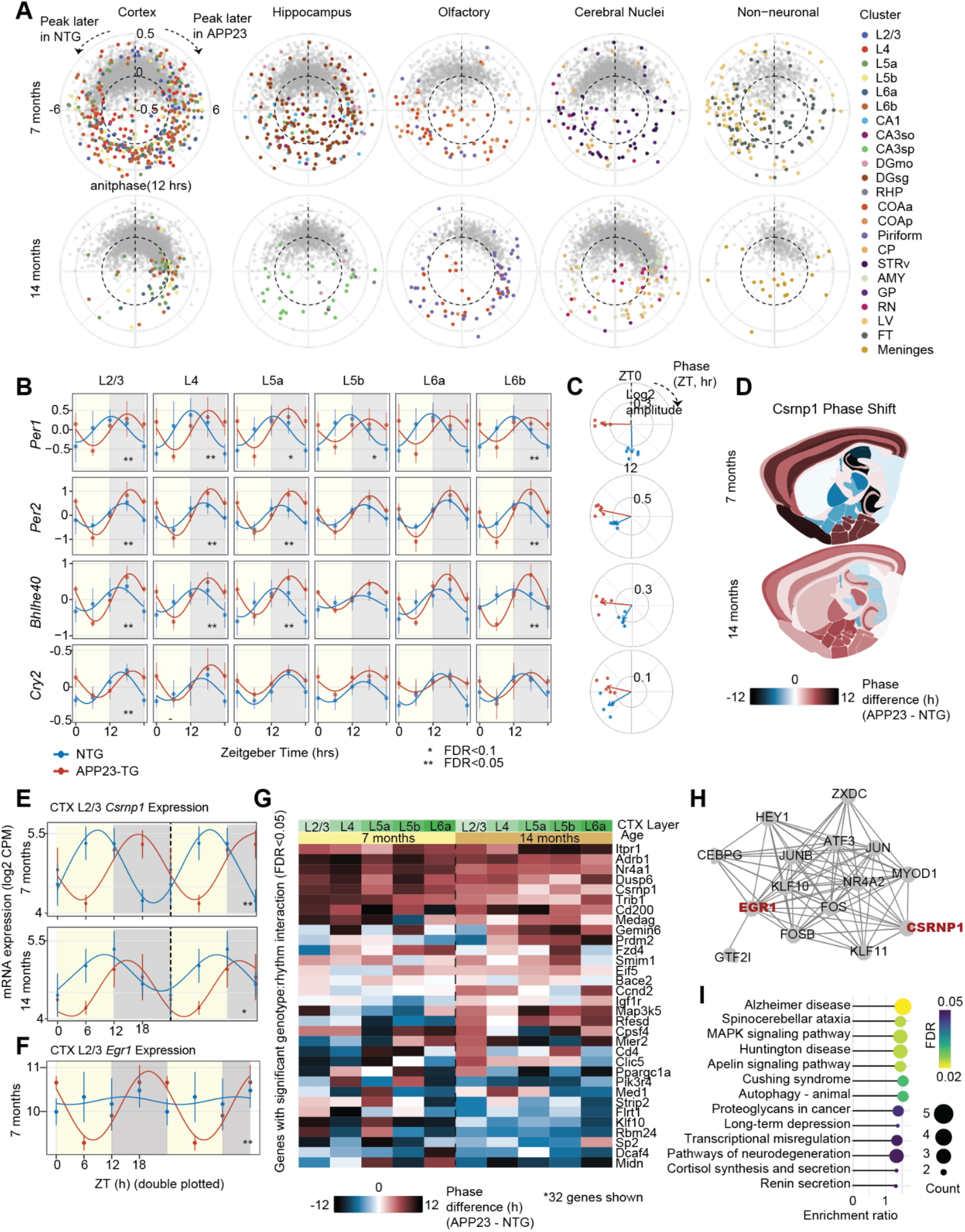
Phase dysregulation of rhythmic gene expression in APP23-TG varies with disease progression. A, Polar plots of DRGs (LRT interaction FDR<0.05) at 7 and 14 months. Radius represents log2 fold change in amplitude (log2 amp APP23 − log2 amp NTG) and angle represents phase difference (θ APP23 − θ NTG, hrs). **B,** mRNA expression and harmonic fits for four differentially rhythmic core clock genes at 7-months across cortical layers (error bars show mean±SD, * p<0.1, ** p<0.05). **C,** Polar plots of phase and log2 amplitude in NTG and APP23-TG mice. Arrows represent mean resultant vectors across cortical layers with vectors colored by genotype. **D,** Spatial map of *Csrnp1* phase difference between APP23-TG and NTG mice projected onto the Allen Brain Atlas at 7 and 14 months. **E,** mRNA expression and harmonic fits for *Csrnp1* in cortex layer 2/3 at 7 and 14 months (error bars show mean±SD, * p<0.1, ** p<0.05). **F,** Same as E for the gene *Egr1* at 7 months**. G**, Phase for genes with significant genotype:rhythm interaction in a full joint model including both ages (LRT FDR<0.1). **H,** Transcription-factor co-regulatory network for the gene set shown in G. Nodes represent TFs whose binding motifs are enriched in DRG promoters (ChEA3); edge thickness reflects the strength of predicted co-regulation. DRGs

The activity-induced transcription factor, *Egr1,* had significantly stronger rhythmicity in APP23-TG compared to NTG at 7 months (Fig. 6F; interaction LRT FDR<0.05) but returned to very low amplitude oscillation in APP23-TG mice at 14 months. Another transcriptional regulator, *Csrnp1* (cysteine-serine-rich nuclear protein 1), stood out for its pronounced phase delay (10 h). *Csrnp1,* which has been linked to genetic risk for AD(*57*), was significantly phase-delayed in cortex layer 2/3 of APP23-TG mice at 7 and at 14 months of age (Fig. 6E; interaction LRT FDR<0.05).

To identify potential drivers of these expression phase shifts, we performed transcription factor motif enrichment analysis on the DRGs using ChEA3(*58*). We found significant enrichment of binding motifs in DRG promoters, including transcription factors *Csrnp1*, *Jun*, *Atf3*, *Fos*, and *Egr1* (Fig. 6H, FDR<0.05, Supplementary Table S11). Interestingly, many of these factors are immediate-early genes that are transcriptionally induced by neuronal activity.

By 14 months, and coinciding with increased amyloid burden, most cortical and hippocampal transcript that gained rhythmicity at 7 months were no longer differentially expressed (Fig. 6A). This finding supports that pre-amyloid changes in rhythmicity may be an adaptive response to early pathological changes, which is later overridden by neurodegeneration as disease advances.

Despite their relative scarcity, DRGs detected at 14 months in APP23-TG mice were functional for pathways directly linked to pathology, including neurodegeneration and cellular stress (FDR<0.05, Fig. 6I, KEGG analysis). Top enriched pathways specifically included Alzheimer’s disease, MAPK signaling, autophagy, and transcriptional misregulation, indicating that DRGs converge on key disease-relevant processes (e.g., protein homeostasis, metabolism, and stress signaling). Together, these findings demonstrate that APP23-TG mice have pronounced, region-specific perturbations in diurnal gene expression timing characterized by phase and amplitude changes in the core molecular clock and relayed to first and second-order output genes that integrate networks fundamental to neurodegeneration.

## Discussion

This study presents a brain-wide, Zeitgeber time (ZT)-resolved spatial transcriptomic atlas of the mouse brain that maps the landscape of brain region-specific diurnal gene expression in health and along the progression of Alzheimer’s pathology. The regulation of 24-hour rhythmicity in the brain is critical for healthy cognition (*2*, *59–62*). Whole-brain microscopy has shown that circadian rhythms are ubiquitous across brain regions(*2*). Transcriptome sequencing further characterized the extent of rhythmic gene expression in cortex and hippocampus(*9*), and single cell RNA-seq of the whole brain confirmed cell type specific regulation of light- vs. dark-phase gene expression(*7*). Our study now provides a spatially resolved, whole-transcriptome map of diurnal gene expression as a foundation for understanding the region-specific regulation of circadian rhythms in health and disease.

Across the brain, our data show that the phase of core-clock gene rhythms are aligned, but the number and composition of rhythmic transcripts differ by region. Subfields of the hippocampal formation and upper cortical layers have the largest number of rhythmic genes(*9*), while subcortical regions and white matter tracts had lower levels of rhythmicity. The phase of peak expression has a bimodal distribution, with most genes peaking several hours before the light/darkness transitions. A notable exception was the thalamic reticular nucleus, a key regulator of sleep(*30*)(*31*), in which several dozen rhythmic genes peak around ZT4. This finding highlights the importance of spatially resolved rhythmic gene expression analysis for a deeper understanding of brain function.

Our ST data reveal differences in rhythmic gene expression along major axes of the cortex and hippocampus. Notably, we found a profound difference in rhythmic gene expression between the caudal visual cortex and rostral sensorimotor areas. Rhythmic expression of core clock components such as *Dbp* was attenuated in caudal cortex, while other genes like *Elk1* had stronger rhythmicity in caudal compared with rostral cortex. A group of neural activity-regulated genes, including *Egr1* and *Egr3*, presented antiphasic expression in caudal vs. rostral cortex.

These results are consistent with a light-driven increase of neural activity in visual cortical areas, whereas motor and somatosensory areas are mainly activated during the dark phase, coinciding with the active phase in mice. In the hippocampus, we confirmed previous reports of large differences in mesor, or mean expression, for hundreds of genes between the dorsal and ventral subfields (*32*, *33*, *35*). Despite these differences in the mesor, diurnal rhythms were largely preserved across the dorsal-ventral axis and we found few differentially rhythmic genes.

Our comprehensive ST data allowed us to analyze alterations in rhythmic transcription in the APP23 model of AD, showing that the brain regions most vulnerable to Aß pathology presented the greatest alterations in the rhythmic transcriptome. Notably, at 7 months of age, before widespread plaque deposition, we observe brain-wide expansion of rhythmic genes and an increase in rhythmic amplitude, especially in cortex. These data extend observations of dysregulated rhythmic gene expression in other models of AD pathology, providing novel spatial context(*9*, *48*, *63*, *64*). In our study, core clock transcripts show phase delays of several hours at 7 months, indicating a shift in molecular timing that parallels circadian behavioral disturbances. The observation of dramatic changes at early disease stages highlight the potential role of circadian dysregulation as a driver or amplifier of subsequent pathological cascades.

Genes that gained rhythmicity in APP23-TG are enriched for activity-regulated immediate-early genes, suggesting neuronal activation may have greater temporal structure in APP23-TG. In cortex, an activity-based gene score was more rhythmic in APP23-TG, supporting this interpretation. Among the IEGs, *Csrnp1* had a robust rhythm with a marked phase delay at seven months that remains detectable, though attenuated, at fourteen months, suggesting that some timing changes persist as disease advances. Importantly, our data do not distinguish gene expression changes which may cause circadian disruption from those which are consequences of altered rhythmicity. Future experiments could test the causal roles of genes highlighted here.

Limitations qualify these conclusions. Our ST data have spatial resolution of ∼50 µm, and each spot includes transcripts from 1-5 or more cells. This can obscure information, for example about genes which oscillate with different phases in different cell types. Single cell sequencing can help to resolve such differences, but previous studies have included a limited number of ZT samples (e.g. 2 time points(*7*)). We designed our study with four ZT samples to allow estimation of both phase and amplitude for diurnal gene expression rhythms across the brain. We focused on a single model of amyloid pathology, APP23-TG, and two ages; extending these studies to tauopathy models, additional ages, and/or human tissue will establish the general validity of our findings. Our data set the stage for these investigations and establish the critical importance of accounting for diurnal rhythmicity in analyses of cell- and region-specific impacts of neurodegeneration.

## Acknowledgments

AI-assisted technology (Grammarly and ChatGTP) were used to improve grammar and reduce text length to fit word count limits.

## Author contributions

Conceptualization: PD

Formal analysis: AG, HR, DB, DC, EAM, PD

Investigation: HR, DW, PD

Visualization: AG, HR, DB, DW

Funding acquisition: PD

Project administration: EAM, PD

Supervision: EAM, PD

Writing – original draft: AG, HR, EAM, PD

Writing – review & editing: AG, HR, DB, DW, EAM, PD

## Data, code, and materials availability

The raw and processed spatial transcriptomics data from this study are available at NCBI accession GSE282203.

## Materials and Methods

### Experimental model details

APP23 transgenic (APP23-TG) mice (B6.Cg-Tg(Thy1-APP)3Somm/J; RRID:IMSR_JAX:030504) and non-transgenic (NTG) littermates control mice were housed in light-tight enclosures at the University of California, San Diego. The mice were given ad libitum food (Teklad rodent diet 8604; Envigo, Indianapolis, IN) and water access. This study used a total of 65 APP23 mice almost equally distributed across sex and genotype. The work presented in this study followed all guidelines and regulations of the UCSD Division of Animal Medicine that are consistent with the Animal Welfare Policy Statements and the recommendations of the Panel on Euthanasia of the American Veterinary Medical Association.

### Experimental design

Prior to beginning any experiments, mice were habituated to a 12h:12h light-dark (LD 12:12) cycle and single housing conditions in custom light-tight cabinets. We then measured diurnal rhythms in locomotor activity (see below). By definition, Zeitgeber time 0 (ZT0) is the time when lights go on and ZT12 is the time when the lights go off. After behavior tests were completed, mice within each genotype and treatment group were randomly assigned for sacrifice at one of four time points: ZT0, ZT6, ZT12, or ZT18. ZT0 and ZT6 tissues were collected in the light; ZT12 and ZT18 were collected in the dark.

### Monitoring of cage locomotor activity

Experimental mice were singly housed in standard cages equipped with IR motion sensors (Actimetrics) and locomotor activity data were acquired using ClockLab Wireless Data Collection and analyzed using ClockLab Analysis 6 (Actimetrics) as previously described(*9*). Mice were entrained to a 12:12 LD cycle. Activity data in LD were collected for at least two weeks. The amount of cage activity (number of IR beam breaks) over a 24 h period was averaged over the recording period and reported in arbitrary units (a.u.)/h. The average number of activity bouts and the average bout lengths were determined for 24 h and for light and dark periods, with new activity bouts defined following an inactivity gap of either 21 min (maximum gap: 21 min; threshold: 3 counts/min) or 1 min (maximum gap: 1 min; threshold: 3 counts/min).

To determine free running period, an independent cohort of mice were placed into constant darkness (DD) for two weeks following LD activity recording. These mice were not used for spatial transcriptomics. Free running period was obtained from the slope of a line fitted to the daily activity onset times during the DD period.

### Tissue collection

Mice were euthanized with CO_2_ followed by decapitation, either in the dark (ZT12 and ZT18) or in the light (ZT0 and ZT6). Brain hemispheres were collected and placed in OCT and then flash frozen in isopentane in liquid nitrogen. One hemibrain from each mouse was cryosectioned at - 18°C sagittally to a thickness of 10 μm (2.8 mm from the midline) using a standard Leica CM1860 cryostat and processed according to the recommended protocols (Tissue optimization: CG000240 Visium 10X Genomics; Gene expression: CG000239). The tissue was immediately mounted on a Visium spatially barcoded slide (10X genomics). The remaining tissue was covered with OCT and kept at -80 °C until it was cryosectioned again starting at the same position to a thickness of 10 μm and mounted onto a Superfrost plus microscope slide (Fisherbrand) for staining. Each section covered approximately 80% of the 5,000 total spots within their fiducial frame. Slides were stored at -80 °C until use.

### Spatial transcriptomics (ST)

Visium spatial gene expression slides and reagents were used according to the manufacturer instructions (10X Genomics). Each capture area was 6.5mm x 6.5 mm and contained 5,000 barcoded spots that were 55 μm in diameter (100 μm center to center between spots), providing an average resolution of about 1 to 10 cells per spot. Optimal permeabilization time was measured at 24 min. Libraries were prepared according to the Visium protocol (10X genomics) and sequenced on a NovaSeq4 (Illumina) at a sequencing depth of 182 million read-pairs by the UCSD genomics gore (IGM). Sequencing was performed with the recommended protocol (read 1: 28 cycles; i7 index read: 10 cycles; i5 index read: 10 cycles; and read 2: 100 cycles), yielding between 175.8 million and 187.9 million sequenced reads. H&E (Hematoxylin, Thermo Cat. No. Thermo; Dako bluing buffer, Dako Cat. No. CS702; Eosin Y, Sigma, Cat. No. 1.09844.1000) staining and image preparation was performed according to the Visium protocol. H&E-stained sections were imaged using a Nanozoomer slide scanner (Hamamatsu) Spatial gene expression assay was performed according to the protocol CG000239. Samples with a sequencing saturation below 50% were sequenced again.

### Pre-processing libraries

Each Visium slice was processed using the SpaceRanger software provided by 10X genomics for mapping reads, aligning spot barcodes to the brightfield tissue image, and generating UMI count table tables. The human APP23 transgene was added to mm10 2020-A reference in order to be able to later quantify transgene expression. As a preliminary quality control measure we analyzed each slice individually using the Seurat ST module(*65*) and inspected the distribution of number of genes detected, number of UMIs, percent mitochondrial reads and number of barcodes passing UMI thresholds. We excluded 4 slices from downstream analysis that had median detected features per spot of less than 500 genes with over 90% sequencing saturation..

Lastly we verified that male and female samples expressed the appropriate sex markers which led us to exclude one more slice due to coexpression of Xist and Y chromosome genes. This yielded 65 high quality, with each combination of experimental covariates having at least 2 replicates except 14 month, non-transgenic males at ZT0 which only had one replicate.

### Integrated Clustering

We used the method PRECAST and performed parameter exploration to find a clustering that best corresponded with the Allen Brain Mouse Reference Atlas [10] both anatomically and transcriptomically. PRECAST allows for integrated clustering across multiple ST datasets while enforcing spatial smoothness and accounting for complex batch effects by computing latent factors with spatially informed priors. Our final clusters were generated by first filtering out spots with less than 500 genes detected and using the following PRECAST parameters: 25 candidate clusters, 25 latent factors, q=25, and 3000 spatially variable genes detected by the SPARK-X package. This yielded 25 final clusters, 1 of which was judged to be of low quality due to a lower mean number of genes detected than other clusters and was removed. 2 clusters corresponded to fiber tracts and spanned anatomical regions such as corpus callosum and medial forebrain bundle system and were merged into one cluster annotated as fiber tracts. This resulted in 23 final anatomically annotated clusters. Marker genes were identified using Seurat’s FindMarkers function and cluster markers were verified to correspond to their anatomical annotation using the Allen Brain Atlas *in situ* hybridization atlas.

### Pseudobulk generation

As a first step to perform cluster-specific rhythmicity analyses we generated pseudobulk gene expression profiles for each cluster and excluded genes that had less than 10 counts in at least one sample. This yielded roughly 12,000 expressed genes per pseudobulk profile. We also excluded samples from any cluster pseudobulk profile that had less than 20,000 total counts or expressed less than 90% of the genes following count thresholding because this indicated the cluster was underrepresented in the sample. After QC 21 of the 23 clusters included at least 28 of the 33 samples. In particular, across all cortical and hippocampal clusters, at least 32 of 33 samples passed these QC thresholds.

### Immunohistochemistry

For immunohistochemistry (IHC), separate age-matched mice were sacrificed and both brain hemispheres were extracted. One hemi-brain was fixed by 4% paraformaldehyde. Fixed APP23 brains were sectioned sagitally at 40 µm using a Leica VT1000S vibratome. Sections were washed three times in PBS, pre-treated with 1% Triton X-100, 10% H2O2 in PBS for 20 min at room temperature, washed again, and incubated for 1 h at room temperature in 10% serum according to secondary antibody species. The sections were incubated with primary antibodies to microglia marker Iba1 (1:500, FUJIFILM Wako Chemicals U.S.A. Corporation, code number 019-19741), neuronal marker NeuN (1:200, MiliporeMillipore Corp., Cat. No. MAB377), amyloid marker 82E1 (1:500, Immuno-biological Laboratories, Cat. No. 10323), and astrocyte marker GFAP (1:200, Invitrogen, Cat. No. PA5-16291) at 4°C overnight. Sections were washed three times, incubated in 1:100 biotinylated secondary antibody (goat anti-rabbit, Vector Laboratories, Cat. No. BA-1000; or horse anti-mouse Vector Laboratories, Cat. No. BA-2000) for 30 min at room temperature, washed again, incubated in biotinylated HRP and avidin (ABC, Vector Laboratories, Cat. No. 30015 and 30016PK-6100) for 1 h in the dark at room temperature and then treated with diaminobenzidine (DAB) Substrate Kit, Peroxidase (Vector Laboratories, Cat. No. SK-4100) for coloration. 20X images were collected using an Olympus VS200 Slide Scanner and analyzed using ImageJ. Cortex GFAP+ area values were normalized to the total cortex area for each mouse.(*9*)

### Plaque analysis

To identify genes that are differentially expressed in plaque-associated spots versus normal spots in the cortex, we used a generalized linear mixed-effects model. Specifically, we labeled all spots as either plaque associated or non-plaque associated, and predicted gene expression with plaque association as a fixed effect and sample as a random effect, to account for cross-sample variability. We performed a likelihood ratio test comparing models with and without plaque fixed effects, to assess significance of coefficients. To estimate the effect of plaque distance on gene expression we calculated the distance of each spot to the nearest plaque-associated spot.

Only spots with a distance to nearest plaque spot less than 600um were included for analysis. We used another mixed effects models to model gene expression as a function of plaque distance, with a random intercept for sample as above. We performed a likelihood ratio test comparing models with and without plaque distance as a fixed effect. Finally, we modeled the relationship between DAM score and plaque distance, with a random intercept for sample. Here, we used a Wald’s test to test for significance of coefficients. For all analyses, plaque FDR correction was used with a cutoff of 0.1.

## Supplementary Text

### Rhythmicity detection and differential rhythmicity analysis

Next we develop a method based on negative binomial regression which tests for rhythmicity while controlling for sex and age (. We model the expression of each gene with a rhythmic component expressed as a sinusoidal function of Zeitgeber time (ZT). This function can be expressed as a linear combination of two Fourier components, which is therefore compatible with a generalized linear model (GLM) framework. To illustrate this let y_t_ represent normalized counts of a gene at ZT=t, let *A* represent the amplitude, 𝜙 represent the phase, and y_0_ represent the mesor. Then our model is as follows:

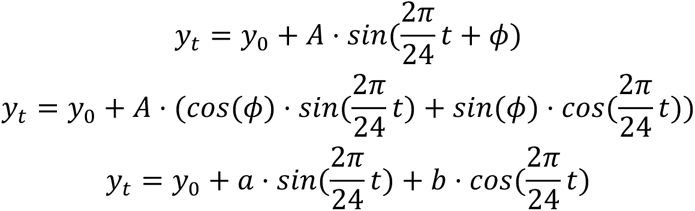

Where a and b represent unknown linear parameter to be fit:

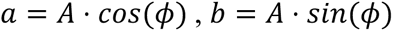

The harmonic model can be tested for significance by removing the ***a*** and ***b*** coefficients and running a likelihood ratio test with the full and reduced model. Once the model is fit estimate amplitude, phase can be recovered via:

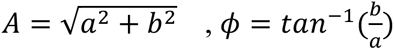

This approach has the benefit of allowing both for the testing of interaction terms directly and fitting age, sex and genotype-specific amplitude and phase all within this same model, unlike standard tools of detecting rhythmicity such as JTK_CYCLE and ARSER. While the approach does not allow for the estimation of period, our experimental approach with four ZT points precludes this possibility. Thus we limit our scope to genes with 24-hour periodicity.

We examined rhythmicity detection results from the glm method described above and found that they were largely consistent with previously established methods implemented in the R package MetaCycle. For genes called as rhythmic by either MetaCycle and/or our approach, there is a high degree of correlation for phase estimates, p-values and relative amplitude estimates. In addition, there is a high degree of overlap in the genes passing chosen FDR thresholds by both methods.

To identify differentially rhythmic genes, we first defined sets of rhythmic genes for each cluster by the method described above in NTG animals and APP23-TG animals separately. For each cluster we then took the union of genes identified in the two genotypes as candidates for genotype-rhythmicity interaction testing. We then ran a likelihood ratio test which included a term modeling the interaction between genotype and the harmonic coefficients and tested it against a reduced model lacking this term. I.e

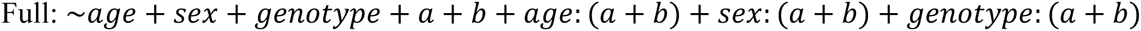

Reduced:

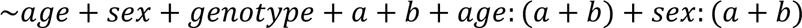

### Permutation-based analysis to control false discovery rate

We used an empirical, permutation-based procedure to ensure that our statistical analysis of differentially rhythmic genes had the expected level of control for false discoveries. We created an empirical null distribution by randomly shuffling the genotype labels of each sample and computing the mean difference in relative amplitude of rhythmic genes. We repeated this shuffling procedure 500 times, and calculated empirical p-values as the fraction of permutations with mean difference greater than that observed in the real data. P-values were adjusted for multiple comparisons across clusters (Fig. 5C).

### Phase density estimation

We took genes identified as rhythmic at an FDR threshold of 0.05 and used the phase estimates to compute a phase density estimation. In brief, kernel density estimation was performed using a Von Mises kernel, and the resulting density estimations were normalized first in the linear sense ( by dividing the density estimate by the sum over all density estimates and the bin width, enforcing a unit area under the estimated density curve) and then for visualization purposes radar plot normalization was performed to normalize the integral of the square of densities in the phase space to 1 (*29*).

### Differential Expression

In order to characterize differential gene expression between APP23-TG and NTG mice across brain regions, we used the same pseudobulk approach and quality control thresholds as input to DESeq2(*25*). For this analysis we reverted to using the Wald test modules of DESeq, as a likelihood ratio test was unnecessary without modeling harmonic effects and it allowed us to compare various combinations of covariates within the same model. After QC thresholding, we tested the effects of genotype, genotype:sex interactions, and genotype:age interactions and visualized the results.

### DAM Score

To understand microglial activation in response to plaque accumulation we devised an approach to compute a disease associated microglia (DAM) score for each cluster in each genotype-age combination. The score is calculated by first taking a list of genes reported to be involved in microglial activation in AD, and subsetted the list to those that were found to be expressed in our data. We then computed the principal components for each cluster-genotype-age combination using the log(CPM+1) values only for those genes. We then plotted the mean score for PC1 for each cluster in spatial coordinates and ran t-tests on the mean DAM scores per cluster and sample between genotypes and ages.

10μm-thick sections that were adjacent to the those used for ST were mounted onto Superfrost plus microscope slides (Fisherbrand) and fixed with 100% methanol for 30 minutes at -20°C. At room temperature, slides were briefly air dried and stained with 0.05% Thioflavin S (Sigma, Cat. No. T1892) in 50% EtOH for 8 minutes followed by three 5-minute washes with 80% EtOH. Sudan black B (0.1% in 70% EtOH) was then applied for 10 minutes, followed by three 5 minute washes with 80% EtOH, one 5 minute was with PBS, and 5 minute was with water. Sections were then coverslipped with Invitrogen ProLong™ Gold Antifade Mounting Medium with DAPI and imaged using an Olympus VS200 Slide Scanner. Due to the difficultly in sectioning the adjacent slide at the same angle after having removed it from the sectioning block, several of the stained sections were cut off at one end. To capture the adjacent tissue across the entire section, multiple 10 μm sections were stained and overlaid. To determine which spots in the Visium slide used for ST were adjacent to Ab plaques, the Thioflavin S-stained sections were overlaid onto the original Visium H&E-stained image. Spots containing or directly in contact with a plaque are defined as plaque associated spots.

### Neural activity score rhythmicity analysis

To estimate neural activity scores, we applied the NeuroEstimator package to pseudobulk profiles in APP23-TG and NTG mice at 7 and 14 months. Neural activity scores were calculated separately for each sample and brain region using default parameters for the package.

To test for rhythmicity, we fit a harmonic regression model, modeling neural activity scores as a sinusoidal function of Zeitgeber time (ZT). A likelihood ratio test (LRT) was performed to compare models with and without harmonic terms. False discovery rate (FDR) correction was applied with a threshold of q < 0.05 to identify significantly rhythmic clusters. Analyses were performed independently for each genotype and age group.

### Pathway enrichment analysis

Pathway enrichment analyses were performed to interpret differentially expressed genes (DEGs) and differentially rhythmic genes (DRGs). Depending on the analysis context, enrichment testing was conducted using either Gene Set Enrichment Analysis (GSEA) or Over Representation Analysis (ORA). GSEA was performed using the WebGestaltR package(*46*) with the KEGG pathway database. ORA was performed using the enrichKEGG function from the clusterProfiler R package(*66*).

---

**Figure S1.**
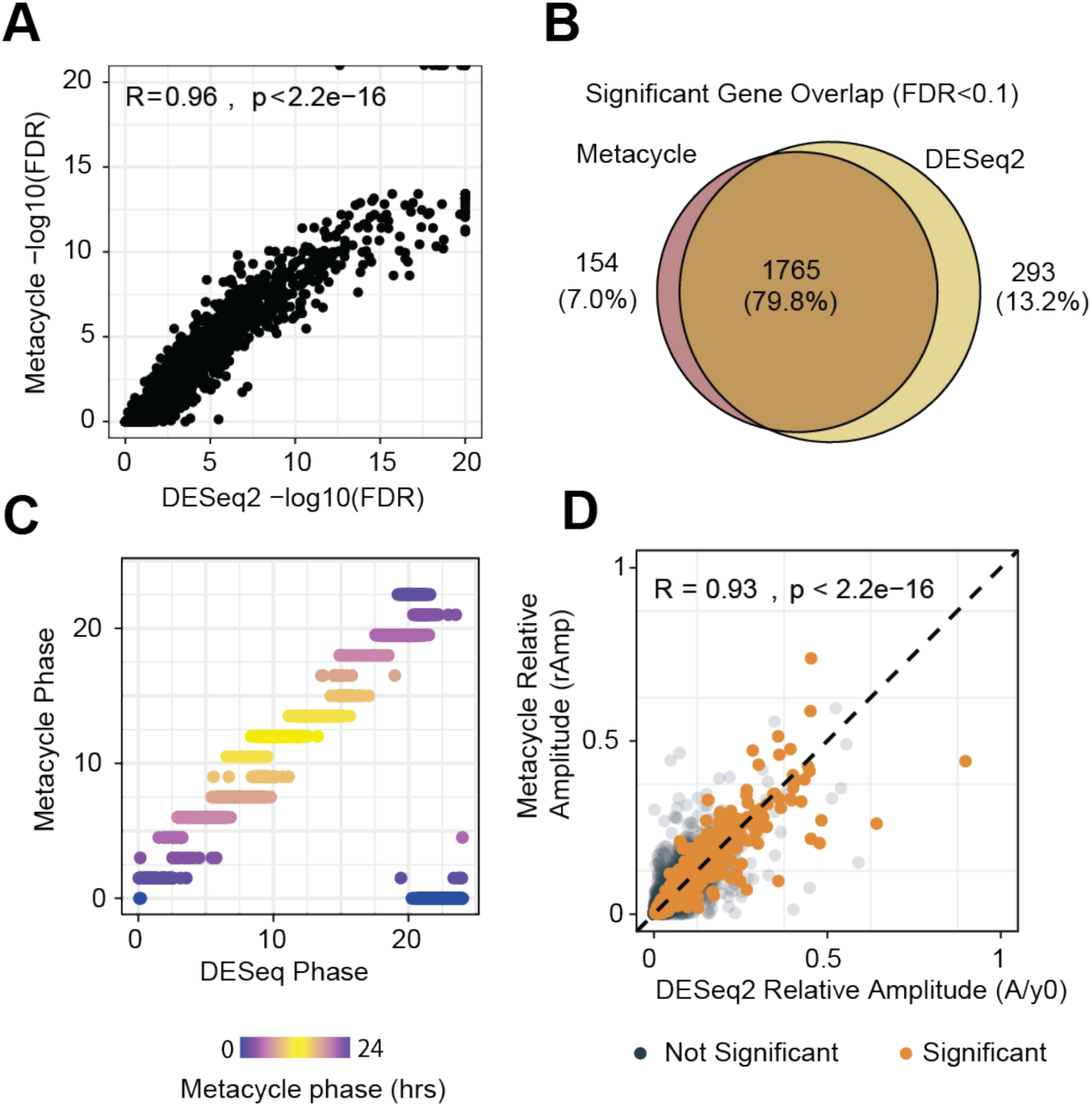
Validation of harmonic regression rhythmicity detection with DESeq2 in cortex layer 2/3 non-transgenic samples. A, Scatterplot of -log10(FDR) values called by MetaCycle versus DESeq2 harmonic regression (spearman correlation=0.96). **B,** Venn diagram of genes passing rhythmicity FDR<0.1 by each method. **C,** Comparison of phase estimates (ZT of peak expression, in hours) from DESeq2 versus MetaCycle phase for significantly rhythmic genes. **D,** Scatter plot of relative amplitude estimates from DESeq2 versus MetaCycle’s meta2d relative amplitude with significantly rhythmic genes (FDR<0.1) highlighted in orange (Spearman correlation=0.93, p<1e-15).

**Figure S2.**
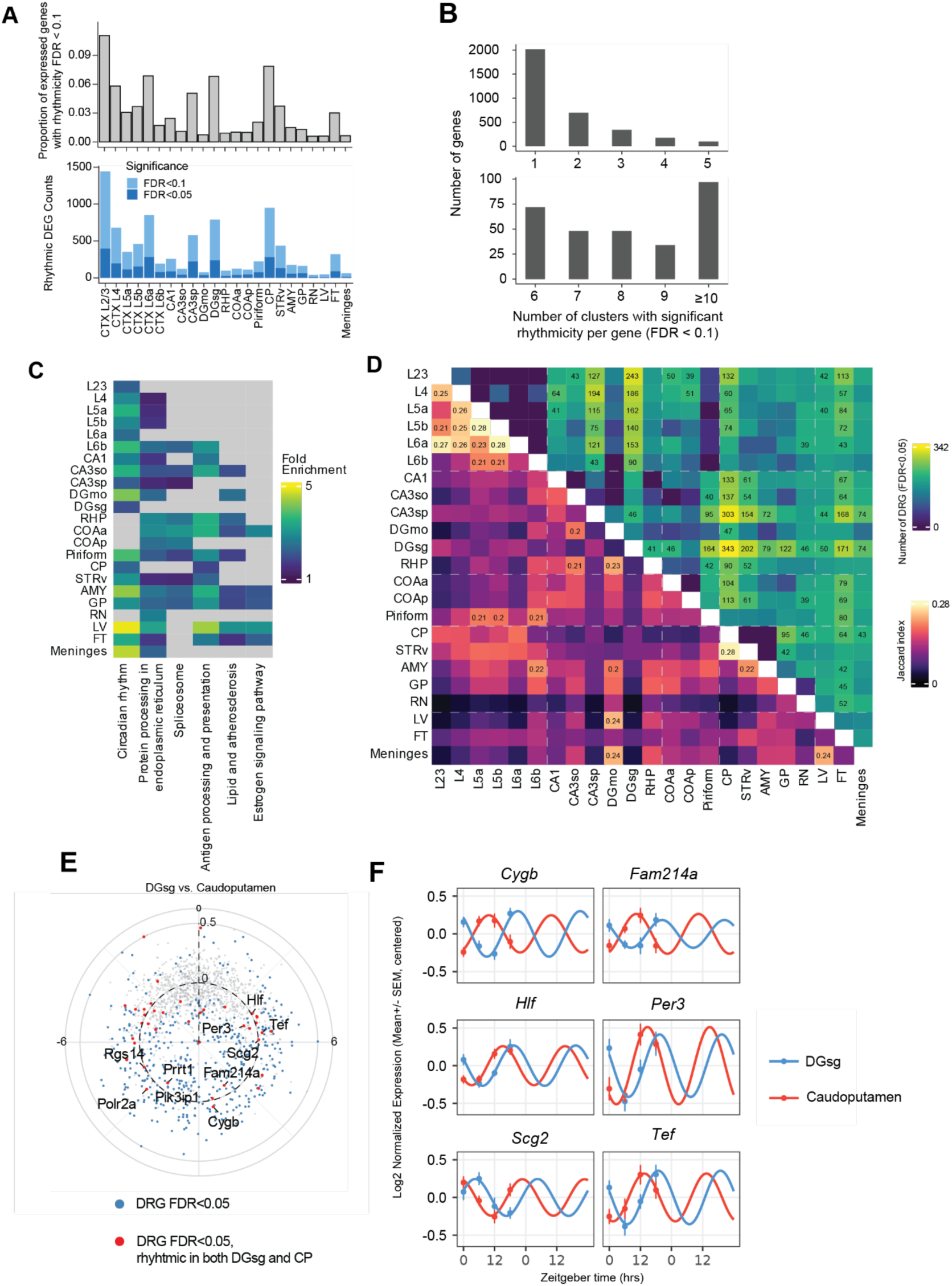
Comparative analysis of gene expression rhythms across brain regions in nontransgenic animals. A, Top, proportion of expressed genes in each cluster with rhythmicity FDR < 0.1. Bottom, number of rhythmic genes, separated by significance threshold (light blue, FDR < 0.1; dark blue, FDR < 0.05). **B,** Number of genes that are significantly rhythmic (FDR < 0.1) in a given number of clusters. The top panel summarizes genes rhythmic in 1-5 clusters, and the bottom panel summarizes genes rhythmic in 6-9 or ≥10 clusters. **C,** Enrichment of KEGG pathways for rhythmic genes (FDR<0.05). Terms found in >1 cluster are shown. Grey cells indicate no significant enrichment (FDR ≥ 0.05). **D,** Overlap of rhythmic genes (bottom left) and number of DRGs (top right, LRT FDR<0.1) for all pairs of clusters. The cluster pairs with the largest overlap or Jaccard index are labeled with the numerical score. **E,** Polar plot of DRGs for an example comparison of two regions, dentate gyrus granule layer (DGsg) versus caudoputamen (CP), with highly distinct rhythmic gene sets. Selected genes with the lowest FDR and similar amplitude are labeled. **F,** mRNA expression and sin fits for DGsg (blue) and CP (red) for DRG. Points (± SEM) show centered log2-normalized expression at each Zeitgeber time, and lines show harmonic regression fits.

**Figure S3.**
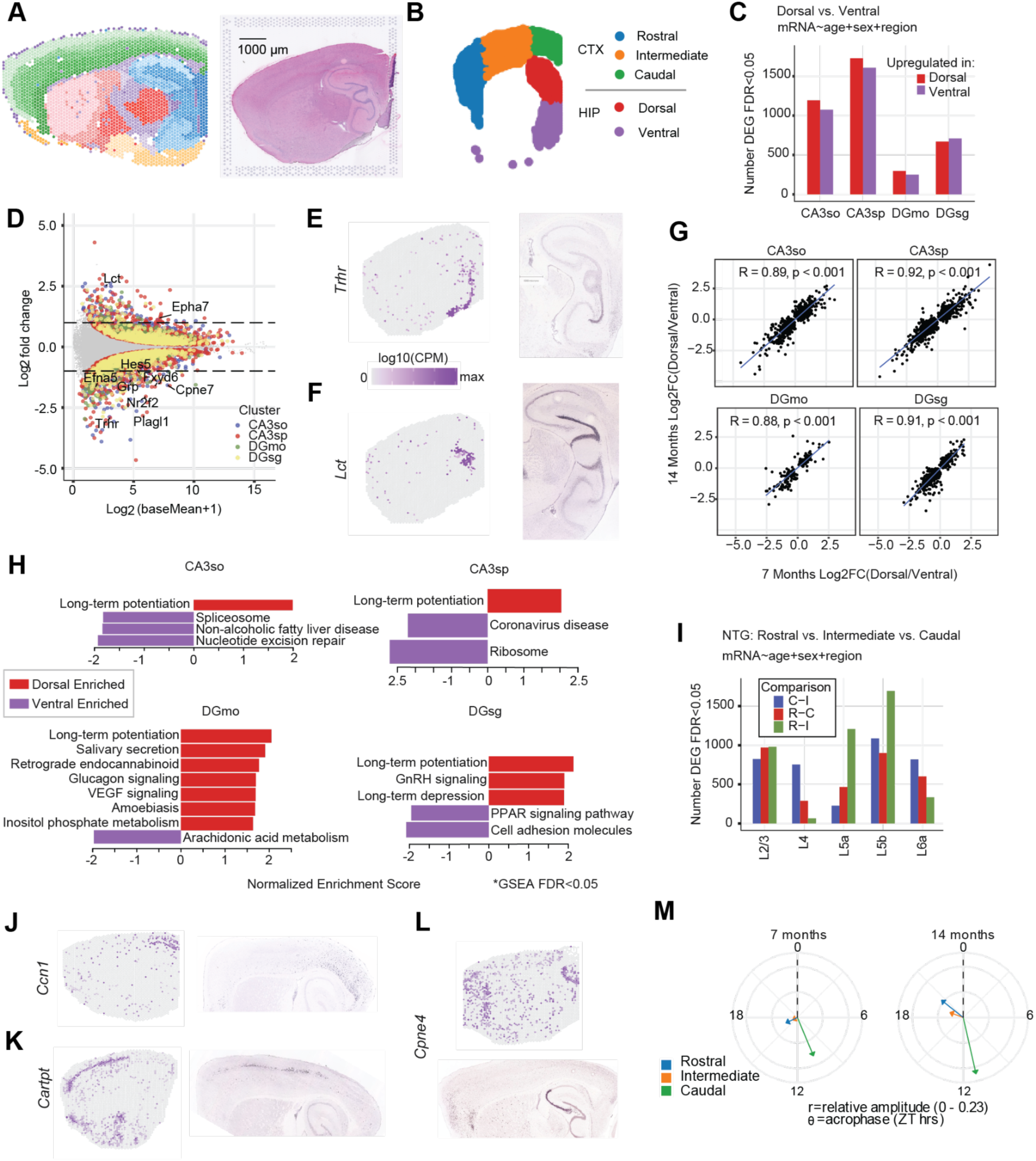
Dorsal versus ventral hippocampal transcriptional differences and relationship to cortical regionalization. A, Sagittal section from the spatial transcriptomic dataset showing clusters overlaid on the brain outline, capturing hippocampus and adjacent rostral, intermediate and caudal cortex. **B,** Annotation of dorsal and ventral hippocampus and rostral, intermediate and caudal cortex used for downstream analyses. **C,** Number of differentially expressed genes (DEGs) between dorsal and ventral hippocampus for each hippocampal cluster (mRNA ∼ age + sex + region, FDR < 0.05). **D,** MA plot of dorsal versus ventral differential expression across hippocampal clusters **E,F,** Spatial expression of dorsal (*Trhr*) and ventral (*Lct*) marker genes in our dataset, and corresponding in situ hybridization images from the Allen Brain Atlas(*23*) (right). **G,** Concordance of dorsal versus ventral log2 fold changes between 7-month and 14-month NTG mice for each hippocampal cluster. **H,** Gene set enrichment analysis of dorsal and ventral log fold changes (GSEA FDR < 0.05). **I,** Number of DEGs identified in pairwise comparisons between rostral, intermediate and caudal cortex across cortical layers (mRNA ∼ age + sex + region, FDR<0.05). **J-L,** Spatial expression of representative cortical regional marker genes and matching Allen Brain Atlas in situ hybridization images. **M,** NeuroEstimator score rhythmic mean resultant vectors for rostral, intermediated and caudal cortex.

**Figure S4:**
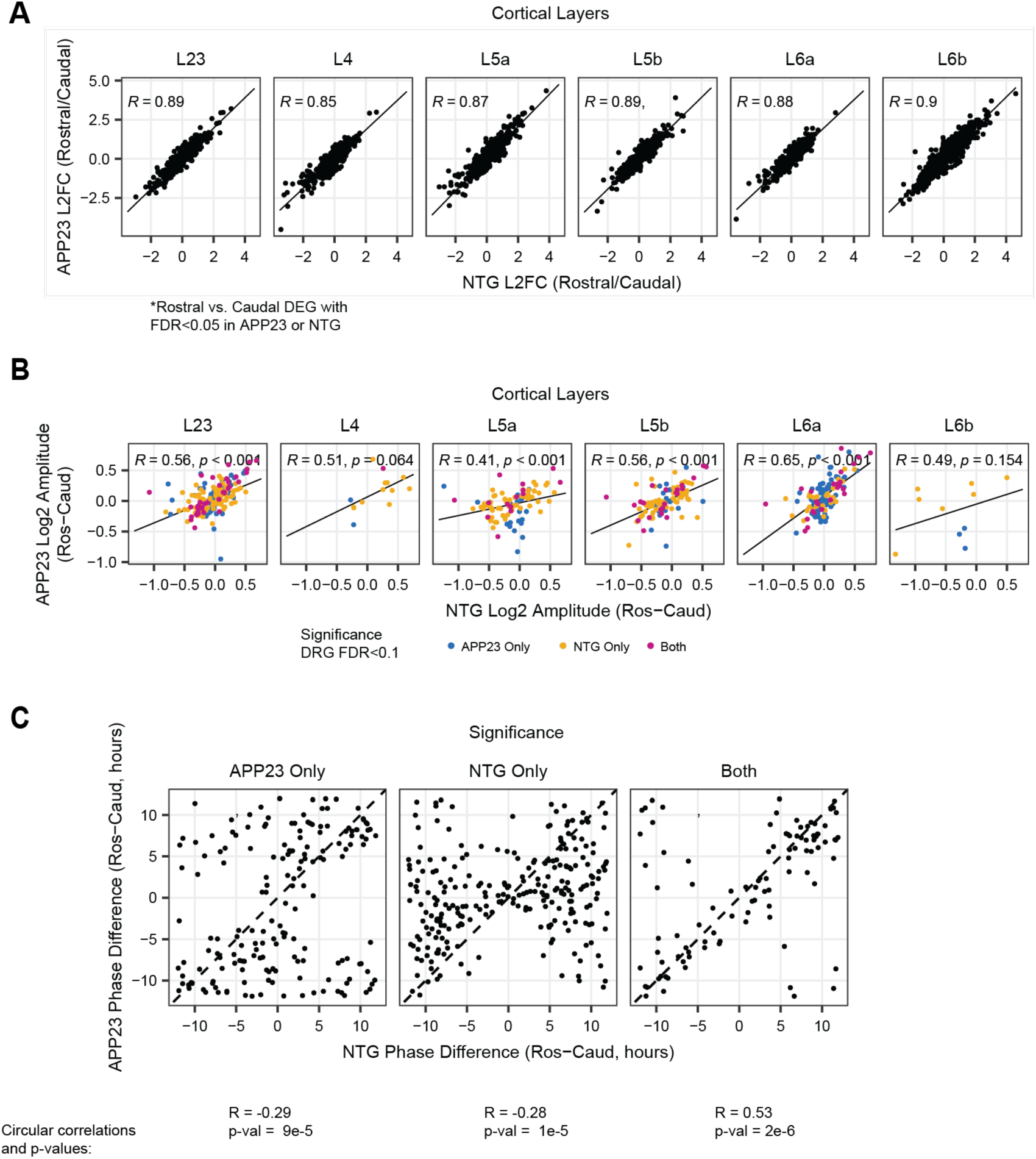
Comparison of differential expression and rhythmicity between rostral and caudal cortex in NTG and APP23. A, Scatterplots of rostral vs. caudal differential expression across cortical layers. Each point is a gene, plotted by its rostral/caudal log2 fold change in NTG (x axis) and APP23 (y axis) **B,** Scatterplots of rostral-caudal rhythmic amplitude differences in NTG (x axis) and APP23 (y axis) for differentially rhythmic genes (FDR< 0.1), colored by rostral-caudal differential rhythmicity significance in NTG only, APP23 only, or both **C,** For all cortical rostral-caudal DRGs (FDR<0.1) scatterplots of phase differences between rostral and caudal in NTG (x axis) and APP23 (y axis), faceted by rostral-caudal differential rhythmicity significance in NTG only, APP23 only, or both. Circular correlations and p values from the Circular R package are shown below(*67*).

**Figure S5:**
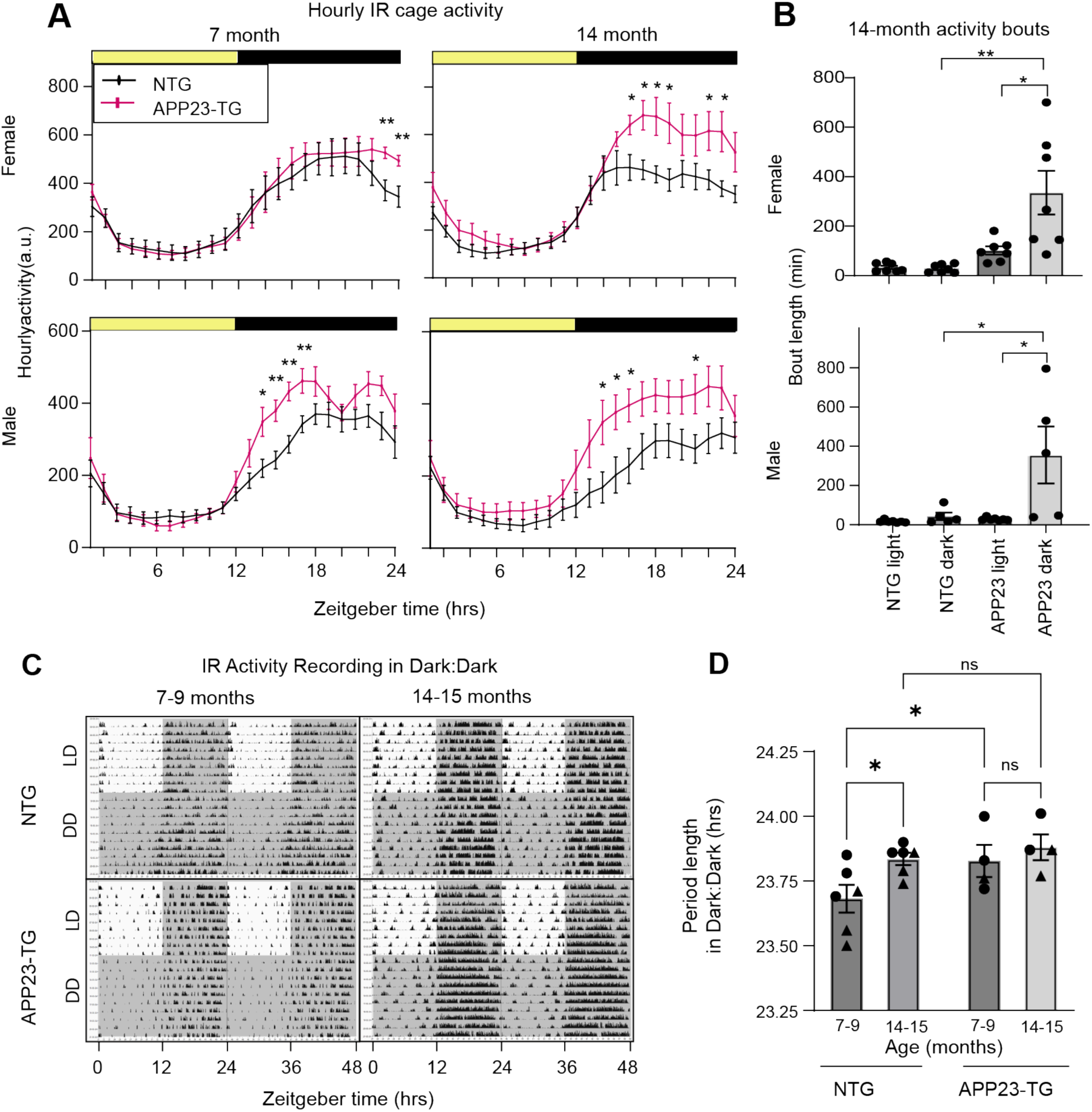
APP23-TG mice exhibit age-dependent alterations in circadian activity and period. A, Mean hourly infrared activity under a 12 h light-12 h dark (LD) cycle at 7 months and 14 months. Mean +/- SEM. Asterisks indicate significant genotype differences at individual Zeitgeber times (ANOVA; * p ≤ 0.05, ** p ≤ 0.01). **B,** Distribution of activity-bout lengths during the light and dark phases measured at 14 months. **C,** Representative actograms of home-cage activity under a 12:12 h light–dark cycle for 7- and 14-month NTG and APP23-TG mice, illustrating hyperactivity during the active period in APP23-TG animals. **D,** Free-running period measured in constant darkness following entertainment at 7 and 14 months of age. The free-running period was significantly longer in APP23-TG compared with NTG mice at 7 months of age (Two-tailed t-test, p ≤ 0.05), but not at 14 months.

**Figure S6:**
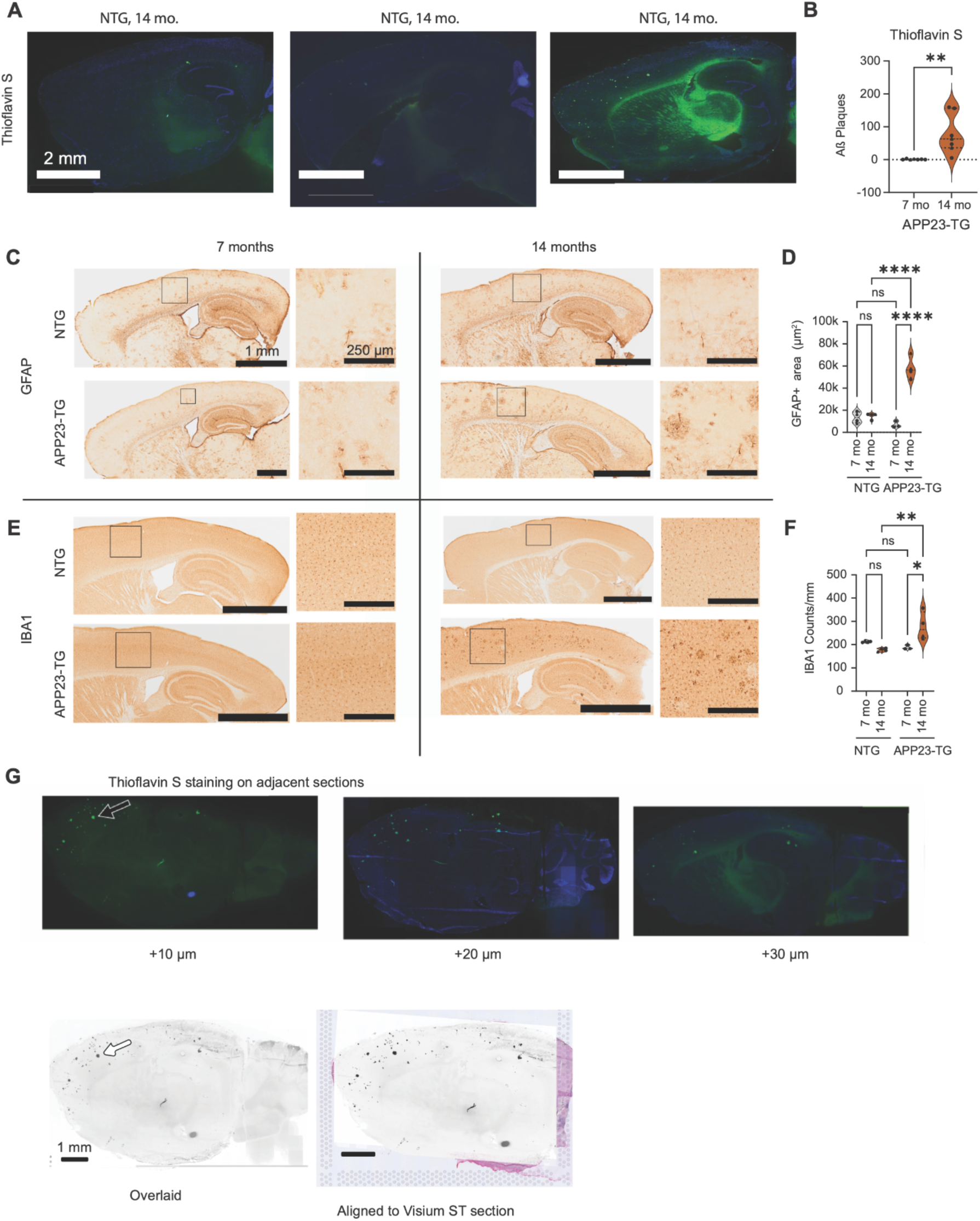
Progressive neuropathology in the APP23 mouse model of AD. A, Thioflavin S staining (green) of Aβ in 14-month old NTG and 7- and 14-month APP23-TG sections (scale bar = 2 mm; DAPI, blue). **B,** Cortical plaque counts in 7- and 14-month APP23-TG sections (n = 7; Two-tailed t test (unpaired); ** p < 0.01) (right). **C,** DAB staining for GFAP (astrocytes) 7- and 14-month NTG and APP23-TG sections. Low-magnification panels (scale bar = 1 mm) with outlines indicate cortical fields zoomed in (scale bar = 250 µm), illustrating progressive astrogliosis. **D,** GFAP positive cortical area in 7- and 14-month NTG and APP23-TG sections (n = 4). **E,** DAB staining for IBA1, illustrating and emerging microgliosis. **F,** Cortical IBA1 counts in 7- and 14-month NTG and APP23-TG sections (n = 4) (bottom right). Circles represent females and triangles represent males (One-way ANOVA with Šídák’s multiple comparison test; ns p > 0.05; * p ≤ 0.05; ** p ≤ 0.01; **** p ≤ 0.0001). **G,** Thioflavin S staining of Aβ in three consecutive, 10 µm sections adjacent to the Visium spatial transcriptomics section. Sequential images were overlaid to reconstruct plaque locations and registered onto the original Visium image (scale bar = 1 mm).

**Figure S7:**
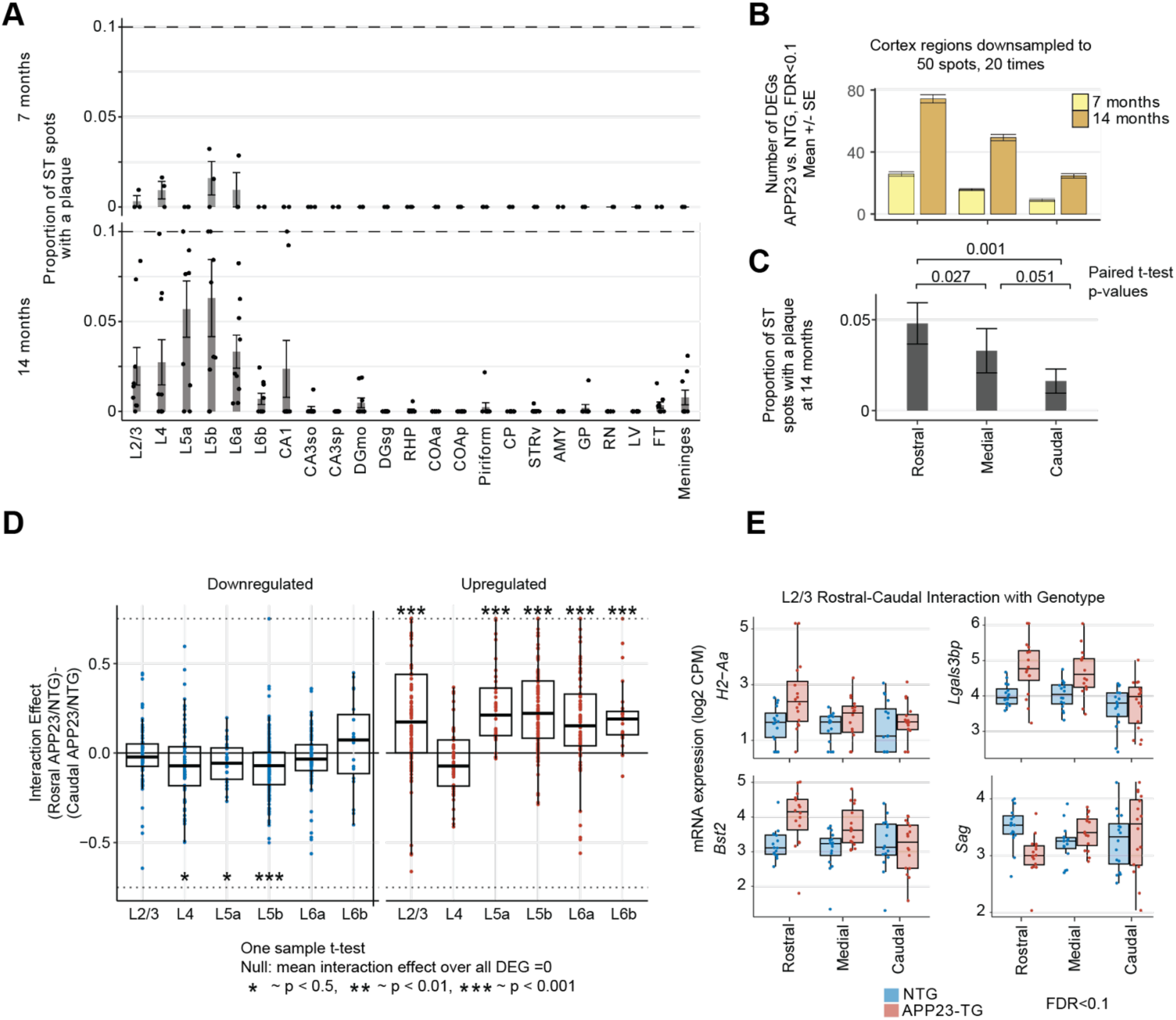
Regional plaque burden, differential expression and interaction effects in APP23-TG cortex. A, Proportion of ST spots containing amyloid plaques in APP23-TG mice at 7 and 14 months (n=16, 9). Bars and error bars show mean ± SE. **B,** Number of DEGs (DESeq2 Wald test FDR < 0.05) in APP23-TG versus NTG across cortical regions, computed on 20 50-spot down-samplings (mean ± SE). **C,** Proportion of plaque-positive spots at 14 months in APP23-TG samples (n=9, bars show mean ± SD); p-values from paired t-tests are shown. **D,** Regional modulation of the genotype effect on expression in each layer, shown separately for genes down-regulated (left) and up-regulated (right) in APP23-TG versus NTG. Each point is the per-gene difference in log₂ fold-change between caudal and rostral cortex (i.e. regional shift of the genotype effect), boxplots summarize the distributions (horizontal line at zero indicates no regional difference), and one-sample t-tests evaluate whether the mean shift differs from zero (*p < 0.05; **p < 0.01; ***p < 0.001). **E,** Boxplots of gene expression for representative genes with significant interaction effects (FDR<0.1) between rostral and caudal cortex. Bars show mean ± SD.

## Supplementary

**Supplementary Table S1: Significantly rhythmic genes in all brain regions in non-transgenic (NTG) animals.** Each tab shows DESeq2 results for one cluster, or the list of shared genes (FDR<0.1 in ≥10 regions). Columns include coefficients for covariates (age_7.months_vs_14.months, sex_M_vs_F) and for the sinusoidal rhythmic components (t_s,t_c); amplitude of the rhythmic component (log2_amplitude, where 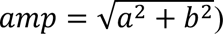); acrophase is the phase of peak expression in ZT hours, 𝜙 = (24/2𝜋) 𝑡𝑎𝑛^−1^(𝑡_(_/𝑡_)_).

**Supplementary Table S2: Significantly rhythmic genes (FDR<0.05, DESeq2) in APP23-TG.** Columns include the coefficients for covariates (age_7_months_vs_14months, sex_M_vs_F) and for the sinusoidal rhythmic components (a,b); phi is the phase of peak expression in radians, 𝜙 = 𝑡𝑎𝑛^−1^ (𝑏/𝑎) 𝑚𝑜𝑑 2𝜋, and phi_hr is the peak phase in ZT (hours), 𝜙_*+_ = 24𝜙/2𝜋; amp is the total amplitude of the rhythmic component 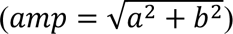.

**Supplementary Table S3: Differentially rhythmic genes (DRGs) between clusters (FDR<0.1).** Tabs are separate analyses for NTG and APP23. Columns include the two regions being compared (test), labels for cluster 1 and cluster 2, test statistics (pvalue, padj from likelihood ratio test comparing models with separate sinusoidal components in the two regions vs. a single shared sinusoidal component), coefficients for covariates (mean cluster 1 vs. cluster 2) and for the sinusoidal rhythmic components for cluster 1, sinusoidal interaction terms for cluster 2, and derived rhythmic parameters per cluster (c1= cluster 1, c2=cluster2) as in table 3. For example, amplitude and phase for cluster 2: 𝑎𝑚𝑝_c2_ = 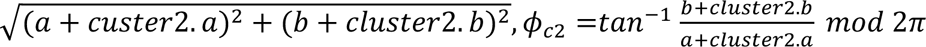

**Supplementary Table S4: Differentially expressed genes (DEGs) in dorsal vs. ventral hippocampus (FDR<0.1).** Each tab includes DEG results for one hippocampal cluster. Columns include gene, baseMean is the mean of DESeq2 size-factor normalized counts, log2FoldChange (dorsal vs. ventral), lfcSE is the standard error of the log2 fold change, stat is the Wald test statistic, p value (Wald test), and FDR_BH (Benjamini–Hochberg false discovery rate).

**Supplementary Table S5: Differentially expressed genes (DEGs) between cortical regions.** Each tab includes DEG results for one cortical layer. Columns include gene, contrast (region comparison e.g., RvI denotes rostral vs intermediate), baseMean is the mean of DESeq2 size-factor normalized counts, log2FoldChange, lfcSE is the standard error of the log2 fold change, stat is the Wald test statistic, p value (Wald test), and FDR_BH (Benjamini–Hochberg false discovery rate).

**Supplementary Table S6: Cortex differentially rhythmic genes (DRGs) in rostral vs. caudal, intermediate vs. caudal, and rostral vs. intermediate regions.** Each tab includes DRG results (FDR<0.1) for one comparison for each cortical cluster in NTG animals. Columns include gene, cluster, rhythmicity FDR in each region (rhythmicity_region_1/2), phase of peak expression in radians and hours (phi_region_1/2, phi_hr_region_1/2), and log2-scale amplitude in each region (amp_region_1/2).

**Supplementary Table S7: KEGG pathway over-representation analysis for rostral-caudal DRGs.** Enrichment analysis was performed for DRGs between rostral and caudal cortex for each layer of the cortex. DRGs with FDR<0.1 were split based on amplitude into rostral- or caudal biased and enrichment analysis was performed using the enrichKEGG function from clusterProfiler (*66*) using layer-specific rhythmic genes as background. Each tab includes results with adjusted p value<0.05 with columns specifying the cortical layer (cluster), KEGG pathway (Description), adjusted p value (p.adjust), and other default outputs from the enrichKegg function.

**Supplementary Table S8: DEGs for APP23 vs NTG per cluster.** Each tab includes DEGs between genotype for all clusters (FDR<0.1) for 7 months and 14 months of age. Columns include cluster, gene, baseMean (DESeq2 size-factor-normalized mean counts), log2FoldChange (APP23/NTG), lfcSE (standard error of the log2 fold change), stat (Wald test statistic), p value, and FDR_BH (Benjamini-Hochberg FDR). Note that Thy1 and humanAPP expression in the APP23 mice reflect the direct effect of the transgene.

**Supplementary Table S9: Differentially expressed genes between Aβ plaque-associated spots and non-plaque associated**. Results from a mixed effects model comparing gene expression in plaque-associated spots with non-plaque associated spots in the cortex (FDR<0.1, LRT: ∼plaque+(1|sample) vs. (1|sample) ). Columns include gene name, p-value, z score, adjusted p-value (padj) and log fold change (lfc).

**Supplementary Table S10: DRGs for APP23 vs NTG per cluster.** Each tab included DRGs with FDR<0.1 at 7 months and 14 months. Columns include cluster, gene, Intercept (APP23 mean), genotype_WT_vs_APP23 (NTG-APP23), sex_M_vs_F, t_s and t_c (APP23 sine and cosine terms), genotypeWT.t_s and genotypeWT.t_c (genotype by time interaction terms), log2-space amplitudes for APP23 and NTG (app_amp, wt_amp) and peak phase in radians and hours for APP23 and NTG (app_phi, app_phi_hr, wt_phi, wt_phi_hr), p value, and fdr_BH (Benjamini-Hochberg adjusted p values). Derived parameters follow the same logic as in Table 3, e.g 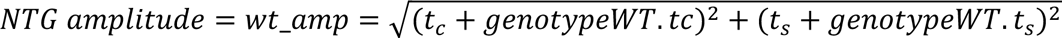

**Supplementary Table S11: ChEA3 transcription factor enrichment for cortical DRGs.** Differentially rhythmic genes (FDR < 0.1) from all cortical clusters (APP23 vs. NTG at 7 months) were submitted to ChEA3 using the rChEA3 R package(*58*). Integrated meanRank and topRank results are filtered to transcription factors that were significant (FDR < 0.05) in at least one underlying ChEA3 library. For each individual library, transcription factors with library-level FDR < 0.05 are shown. For integrated results columns report transcription factor rank within the library (Rank), transcription factor symbol (TF), ChEA3 enrichment score (Score), the contributing ChEA3 library (Library), and the list of input genes overlapping each transcription factor target set (Overlapping_Genes). For individual libraries, column additionally report the number of overlapping genes between the input set and target set (Intersect), total size of the target set (Set length), Fisher’s Exact Test p-value (FET p-value), library-level FDR, and the corresponding odds ratio. (FDR<0.05) are highlighted red. **I,** Bubble plot of KEGG pathway enrichment of DRGs (APP23-TG vs. NTG).

